# Extremely reduced supergroup F *Wolbachia*: transition to obligate insect symbionts

**DOI:** 10.1101/2021.10.15.464041

**Authors:** Sazzad Mahmood, Eva Nováková, Jana Martinů, Oldřich Sychra, Václav Hypša

## Abstract

*Wolbachia* are widely distributed symbionts among invertebrates that manifest by a broad spectrum of lifestyles from parasitism to mutualism. *Wolbachia* Supergroup F is considered a particularly interesting group which gave rise to symbionts of both arthropods and nematodes, and some of its members are obligate mutualists. Further investigations on evolutionary transitions in symbiosis have been hampered by a lack of genomic data for Supergroup F members. In this study, we present genomic data for five new supergroup F *Wolbachia* strains associated with four chewing lice species. These new strains in different evolutionary stages show genomic characteristics well-illustrating the evolutionary trajectory which symbiotic bacteria experience during their transition to mutualism. Three of the strains have not yet progressed with the transition, the other two show typical signs of ongoing gene deactivation and removal (genome size, coding density, low number of pseudogenes). Particularly, *w*Meur1, a symbiont fixed in all *Menacanthus eurysternus* populations across four continents, possesses a highly reduced genome of 733,850 bp with a horizontally acquired capacity for pantothenate synthesis. Comparing with other strains showed *w*Meur1 genome as the smallest currently known among all *Wolbachia* and the first example of *Wolbachia* which has completed genomic streamlining known from the gammaproteobacterial obligate symbionts.

## Introduction

*Wolbachia* present a unique example of diversity and phenotypic flexibility found within a single monophyletic group of bacterial symbionts. Originally described as a causative agent of cytoplasmic incompatibility in *Culex pipiens* mosquitoes, the genus is today known to be widely distributed among many arthropods and some nematode species (1). The diversity of their lifestyles spans from parasites to obligate mutualists. Phylogenetically, *Wolbachia* form several distinct clusters, usually called supergroups (2) which have developed their own characteristic features and tendencies. For example, while most supergroups seem to be specific to arthropods, few are reported exclusively from filarial nematodes. A particularly interesting group is supergroup F, the only supergroup known to infect both arthropods and nematodes (2-4). Moreover, some members of this supergroup are highly adapted strains with degraded genomes, which can maintain a mutualistic relationship with their hosts (5, 6).

The rapidly growing number of *Wolbachia* genome assemblies now allows for evolutionary and functional comparisons and identification of the characteristics underlying different life strategies (4, 7). However, the distribution of the available data across the supergroups and host taxa is extremely uneven and biased. The recent meta-analysis performed by Scholz et al. (1) included an impressive number of 1,166 *Wolbachia* genomes or genome drafts, but the majority of them (1,018) originated from dipterans, almost exclusively from *Drosophila* (1,011). Similarly, regarding the taxonomic diversity of *Wolbachia*, 1,055 of the genomes represented supergroup A, while only 11 belonged to supergroup F and originated from three different hosts. This uneven distribution of genomes is likely to reflect the difference in attention paid to model and non-model organisms, rather than the real diversity of *Wolbachia*. Occasional screenings suggest that the supergroups underrepresented by genomic data may encompass a high diversity of *Wolbachia* strains. As an example, supergroup F, currently represented by four genomes and genome drafts from two nematodes and two arthropods (1, 8-10), seems to contain a wide variety of *Wolbachia* from different hosts when screened for specific *Wolbachia* genes (2, 11-13).

Phthiraptera belong to the insect taxa which have been screened specifically for the presence of *Wolbachia* symbionts and seem to be frequently infected. Kyei-Poku et al. (14) performed a PCR-based screening of 19 species, encompassing both sucking lice of the suborder Anoplura as well as the chewing lice of the suborders Amblycera and Ischnocera. They showed that all the tested samples produced specific *Wolbachia* markers, in some cases suggesting the occurrence of multiple strains. Since the screening was based on specific phylogenetic markers, genomic data is not available for these symbionts. It is therefore difficult to hypothesize on the nature of these symbiotic relationships and role of these *Wolbachia* for the hosts. From Anoplura that feed exclusively on vertebrate blood, several obligate symbionts of different phylogenetic origins have been characterized, and for some, their role in provisioning B vitamins has been demonstrated (15-21). Of the sucking lice included in the Kyei-Poku et al. (14) screening, this is the case for *Riesia* in *Pediculus* and *Phthirus* (16, 22), and *Legionella polyplacis* in *Polyplax serrata* (20, 21). Since the mutualistic role of these symbionts is well established, it is likely that *Wolbachia* do not play a nutritional role in these lice. They may rather be accompanying commensals or even parasites, as shown in many other insects. In chewing lice, the situation is less clear. These ectoparasitic insects are likely not a monophyletic group (23) and their feeding strategies are more diverse than in Anoplura. While all chewing lice are ectoparasites living in the fur or feathers of their hosts, the source of food varies among the groups, and in several cases their diet may also include host’s blood (24, 25). Currently, a single genome is available for a chewing louse symbiont (26). This symbiont, described from slender pigeon louse *Columbicola wolffhuegeli*, is phylogenetically related to the genus *Sodalis* within Gammaproteobacteria. Its genomic characterization revealed features resembling other obligate symbionts in insects, namely strong size reduction and shift of GC content (797,418 bp, 31.4% of GC), but did not provide any clear evidence for its function in the host (26).

In this study we investigate the nature of *Wolbachia* symbionts in several species of chewing lice. We focus primarily on the widely distributed species *Menacanthus eurysternus*. This chewing louse has cosmopolitan distribution (27) and is known to parasitize a broad spectrum of passeriform and several piciform bird species (24). This allows us to test the obligate nature of *M. eurysternus*-associated *Wolbachia* across a broad range of samples. To assess the genomic characteristics and capacity of the symbiont, we assemble metagenomic data from *M. eurysternus* and reconstruct the complete genome of its *Wolbachia* symbiont. Finally, for comparative reasons, we assemble additional genome drafts from several chewing lice samples with metagenomic data available in SRA database (28).

## Methods

### Material and DNA extraction

Samples of *Menacanthus eurysternus* were collected across a large geographic distribution (Supplementary data 1) from 2000 to 2016. For 16S rRNA gene amplicon analysis, DNA templates were extracted from 54 individuals using QIAamp DNA Micro Kit (Qiagen) (Supplementary data 1). DNA template for metagenomics was isolated from pool of 8 individuals collected from one specimen of *Fringilla coelebs morelatti* GA72. To avoid environmental DNA contamination, lice were washed with pure ethanol (3x for 30 min) in Mini-rotator (Bio RS-24) and DNA was extracted with QIAamp DNA Micro Kit (Qiagen). Concentration of the isolate was quantified with Qubit 2.0 Fluorometer (Invitrogen, Carlsbad, CA, USA) and the integrity of DNA was verified on agarose gel electrophoresis (1,5%). NEBNext® Microbiome DNA Enrichment Kit (New England BioLabs) was used for increasing the proportion of bacterial DNA (via the procedure of selective binding and disposing of methylated host DNA). Final DNA concentration was quantified with a Qubit 2.0 Fluorometer using High Sensitivity reagents.

### 16S rRNA gene amplicon sequencing and analysis

The diversity and distribution of microbial associates in *Menacanthus eurysternus* samples were assessed using a 16S rRNA gene amplicon sequencing protocol developed by our group (29). Briefly, multiplexing was based on a double barcoding strategy using fused primers with 12-bp Golay barcodes in forward primer 515F, and 5-bp barcodes within the reverse primer 926R (30, 31). An 18S rRNA gene blocking primer (29) was involved in all PCR reactions to ensure sufficient yields of 16S rRNA gene amplicons from the metagenomic templates.

*M. eurysternus* samples were part of a highly multiplexed library containing 384 samples altogether. In order to control for amplification bias and contamination, two positive controls (commercially purchased mock communities ATCC® MSA-1000™ and ATCC® MSA-1001™), and two negative controls for PCR amplification were processed along with *M. eurysternus* samples (complete metadata including barcodes are available in Supplementary data 1. The purified library was sequenced on Illumina Miseq using V2 chemistry with 500 cycles (Norwegian High Throughput Sequencing Centre, Department of Medical Genetics, Oslo University Hospital).

The raw fastq data were processed into the OTU table with an in-house workflow combining Usearch (32) and Qiime 1.9 (33) scripts as described previously (29). Taxonomic classification was assigned to individual OTUs using BLAST searches of representative sequences against the SILVA 138 database (as of February 2021). Non-bacterial OTUs and potential contaminants found in the negative controls were cleaned from the data via a series of decontamination processes using different levels of stringency to evaluate the overall pattern of *Wolbachia* dominance and ubiquity in *M. eurysternus* microbiomes. While the less stringent decontamination involved eliminating 12 OTUs shared by both negative controls, the strict decontamination removed every OTU found the negative controls (35 OTUs altogether). The details on the control profiles and eliminated OTUs can be found in Supplementary data 1. The decontaminated datasets were rarefied in 5 iterations at a level of 1000 and 2000 reads and imported into RStudio (34) using phyloseq package (35). Compositional heat maps were produced for the 20 most abundant OTUs and ordered to reflect the phylogenetic relationship among analyzed *M. eurysternus* samples (complete COI phylogeny available in Supplementary figure 1).

### *Metagenomic sequencing and assembly (*Menacanthus eurysternus)

The shotgun genomic libraries were prepared from the enriched gDNA of *M. eurysternus* GA72 sample using the Hyper Library construction kit from Kapa Biosystems (Roche). The library was sequenced in a multiplexed mode on one SP lane of NovaSeq 6000 for 251 cycles from both ends of the fragments. Fastq files were generated and demultiplexed with the bcl2fastq v2.20 Conversion Software (Illumina). The quality of 145,910,396 paired reads was checked with FASTQC and the data were trimmed by the Bbduk tool (36) to a minimal phred score of 20. Spades with the option --meta was used to assemble the metagenome. Initially, *Wolbachia* contigs were identified by blastn searches (37) using the complete set of genes from the *Cimex lectularius* symbiont wCle (ACC) as a query. *Wolbachia* origin of the preselected contigs was verified by blastn searches against the NCBI nt database. The amplicon analyses indicated the presence of two different *Wolbachia* strains. Based on the considerable length difference between the first contig and the rest of *Wolbachia* contigs (732,857 bp vs 26,534 and less) and different GC contents (28% and >33%), we hypothesized that the first contig may be almost complete genome of one strain, while the others represent a second strain. To test this possibility by closing the genome of the first *Wolbachia* strain, we used two approaches. First, we extended the longest contig by aTRAM 2.0 (38) and closed it into a 733,850 bp long circular sequence. Second, using specific primers designed based on the longest contig, we sequenced the missing part and completed the genome into a circular sequence identical with the result from aTRAM based approach. The remaining 189 contigs were considered as parts of the second strain of *Wolbachia*.

### Screening and assembly of chewing lice SRA

To check for presence of *Wolbachia* in other chewing lice, we screened the available metagenomic data in SRA (28) (Supplementary table 1). Assembling of the reads and detection of *Wolbachia* were done in the same way as for *M. eurysternus* (described above). To identify candidate *Wolbachia* contigs, we used two genomes as queries, *w*Cle and the newly assembled complete genome from *M. eurysternus*. Of five assemblies in which *Wolbachia* contigs were identified, two contained only a few short contigs with low coverage and were not included in the subsequent analyses (Supplementary table 1). For the remaining three assemblies we extracted *Wolbachia* genome drafts with different degree of fragmentations (from 9 contigs in *Meromenopon meropis* to 386 in *Alcedoecus sp*.). Since their sizes did not deviate from the common size of the other *Wolbachia* genomes (Table 1) and the completeness assessed by BUSCO was also comparable to other *Wolbachia* (Supplementary table 2; see below for BUSCO analyzes), we considered these sets of contigs as representative genome drafts.

**Table 1:** Comparison of the main genomic characteristics of the nine *Wolbachia* strains from supergroup F.

| strain | host | genome size | proteins/hypothetical | %GC | busco Rickettsiales/Proteobacteria | transposases | phage-related phages/RAST | mobile elements | ankyrin repeats | intergenic max. | intergenic average | pseudogenes | coding density |
| --- | --- | --- | --- | --- | --- | --- | --- | --- | --- | --- | --- | --- | --- |
| wMeur1 | *Menacanthus eurysternus* | 733,850 | 592/72 | 28 | 85.7/68.5 | 0 | 0/0 | 0 | 0 | 2095 | 370 | 17 | 76 |
| wMmer | *Meromenopon meropis* | 1,005,754 | 734/142 | 29 | 91.8/73.5 | 0 | 0/0 | 0 | 1 | 3771 | 680 | 32 | 66 |
| wMeur2 | *Menacanthus eurysternus* | 1,015,603 | 1,032/306 | 36 | 98.6/83.1 | 1 | 0/9 | 6 | 9 | 2055 | 171 | 85 | 84 |
| wPaur | *Penenirmus auritus* | 1,198,730 | 1,202/395 | 36 | 97.0/82.2 | 3 | 19/23 | 13 | 19 | 2247 | 208 | 144 | 82 |
| wAlce | *Alcedoecus* sp. | 1,479,761 | 1,506/470 | 36 | 97.3/79.9 | 8 | 88/17 | 21 | 9 | 2355 | 183 | 145 | 80 |
| wMelo | *Melophagus ovinus* | 1,656,288 | 1639/520 | 36 | 98.6/82.7 | 4 | 21/78 | 21 | 30 | 2483 | 169 | 137 | 78 |
| wCle | *Cimex lectularius* | 1,250,060 | 1,454/595 | 36 | 97.8/82.2 | 0 | 56/14 | 68 | 26 | 1522 | 239 | 459 | 81 |
| wOc | *Osmia caerulescens* | 1,232,261 | 1,239/424 | 36 | 98.7/83.5 | 1 | 16/37 | 12 | 19 | 1907 | 172 | 167 | 83 |
| wMhi | *Madathamugadia hiepei* | 1,025,329 | 1,081/372 | 36 | 93.1/78.6 | 1 | 0/5 | 23 | 8 | 2450 | 207 | 149 | 79 |

Completeness assessment and annotations were done for both *M. eurysternus* strains and the three SRA-derived strains by the same procedure. Completeness was assessed in BUSCO v5.1.2 with two different references, rickettsiales_odb10.2019-04-24 and proteobacteria_odb10.2019-04-24. Functional annotations of the genes were obtained by RAST (39). The presence of phage-related sequences was further checked by PHASTER (40). Potential pseudogenes were identified by Pseudofinder (41) based on annotation obtained from Prokka (42). Possible horizontal gene transfers (HGT) were identified by diamond blastx (43) against NCBI nr database with complete set of annotated genes as queries, e-value set to 10 and number of hits to five. Assignment of the genes to clusters of orthologous groups (COGs) was done in web-based eggNOG-Mapper (44). To visualize sharing of the genes across the supergroup F strains, we plotted results of the orthofinder analysis by UpSetR package of R (45).

### Phylogeny

To determine the position of the new strains, we designed two different matrices. First, since our preliminary analyses suggested that the new strains belonged to supergroup F, the “*fbpA_coxA*” nucleotide matrix was built to represent this supergroup. The matrix contained 48 F strains, and 23 additional strains representing other supergroups. The genes were retrieved from NCBI (https://www.ncbi.nlm.nih.gov/), pubMLST (46) and our new assemblies (Supplementary table 3). Alignment was done in MUSCLE (47). The web-based IQ-TREE tool (48) was used to select the best models (TN+F+I+G4 and TVM+F+G4 models for *fbpA* and *coxA* respectively) and to perform phylogenic analysis. To verify the position of the new strains within supergroup F, we designed an amino acid “*multigene matrix*”, restricted to the strains for which genomic data are available. This set contained all available genomes for supergroup F and several additional genomes representing other major supergroups (Supplementary table 3). For the included genomes, we identified 101 shared single copy orthologs by Orthofinder (49), aligned them in MUSCLE (47) and removed the unreliably aligned sites by Gblocks (50). The final concatenated matrix containing 23,218 positions was analyzed by two different approaches. Maximum likelihood analysis was done in IQ-TREE with HIVb+F+I+G4 selected as best model. Since our data contained several long-branch sequences we used in addition PhylobayesMPI (51) with CAT-GTR model to minimize possible artifacts (52). This analysis was run for 50,000 generations under two different coding systems, first coding for each amino acid, second with amino acids recoded by the Dayhoff6 system.

### Genomic and metabolic comparisons

Genomic analyses and comparisons were done for the nine strains of supergroup F for which complete genomes or drafts are available (Supplementary table 3). Average nucleotide diversity (ANI) was calculated using a web-based ANI calculator (53). Synteny of the genomes was analyzed in Mauve (54) implemented in Geneious (55). Assessment of metabolic capacities was done using the web-based tools Blastkoala and KEEG mapper (56). To obtain a metabolic overview comparable with other *Wolbachia* supergroups, we adopted the scheme used by Lefoulon et al. (10) and extended its content with comparison of amino acids synthesis.

## Results

### *Amplicon screening of* M. eurysternus

On average, 16S rRNA gene sequencing yielded 6,686 reads per sample under less stringent decontamination and 5,911 reads under the strict decontamination (see Materials and methods). The mock communities yielded, on average, 20,876 reads for the equally composed samples and 48,356 for the staggered communities. We were able to recover the expected profiles for both equal and staggered DNA template, including overrepresentation of *Staphylococcus epidermidis* (ATCC 12228) and vast underrepresentation of *Rhodobacter sphaeroides* (ATCC 17029) reported previously by the manufacturer (Supplementary data 1). Within the staggered communities, we retrieved all three extremely low abundant taxa (0.04%, Supplementary data 1). The presence of an eleventh OTU of the genus *Granulicatella* however pointed out marginal (tens of reads) well-to-well contamination between positive and negative controls (details in Supplementary data 1). In all the datasets decontaminated under different stringency (see Materials and methods and Supplementary figure 2), *Wolbachia* OTUs clearly dominate *M. eurysternus* microbiomes. While the analyzed individuals generally associate with a single *Wolbachia* (OTU2), some show a dual *Wolbachia* infection. However, the number of *Wolbachia* OTUs does not correlate with either geographic or phylogenetic origin of the analyzed hosts (Figure 1). Although few individuals in Figure 1 seemingly lack *Wolbachia*, these samples did not meet the rarefaction thresholds of 1000 and 2000 reads in the strictly decontaminated dataset (Supplementary figure 2) thus confirming a robust pattern of *Wolbachia* ubiquity across diversified populations of *M. eurysternus*.

**Figure 1:**
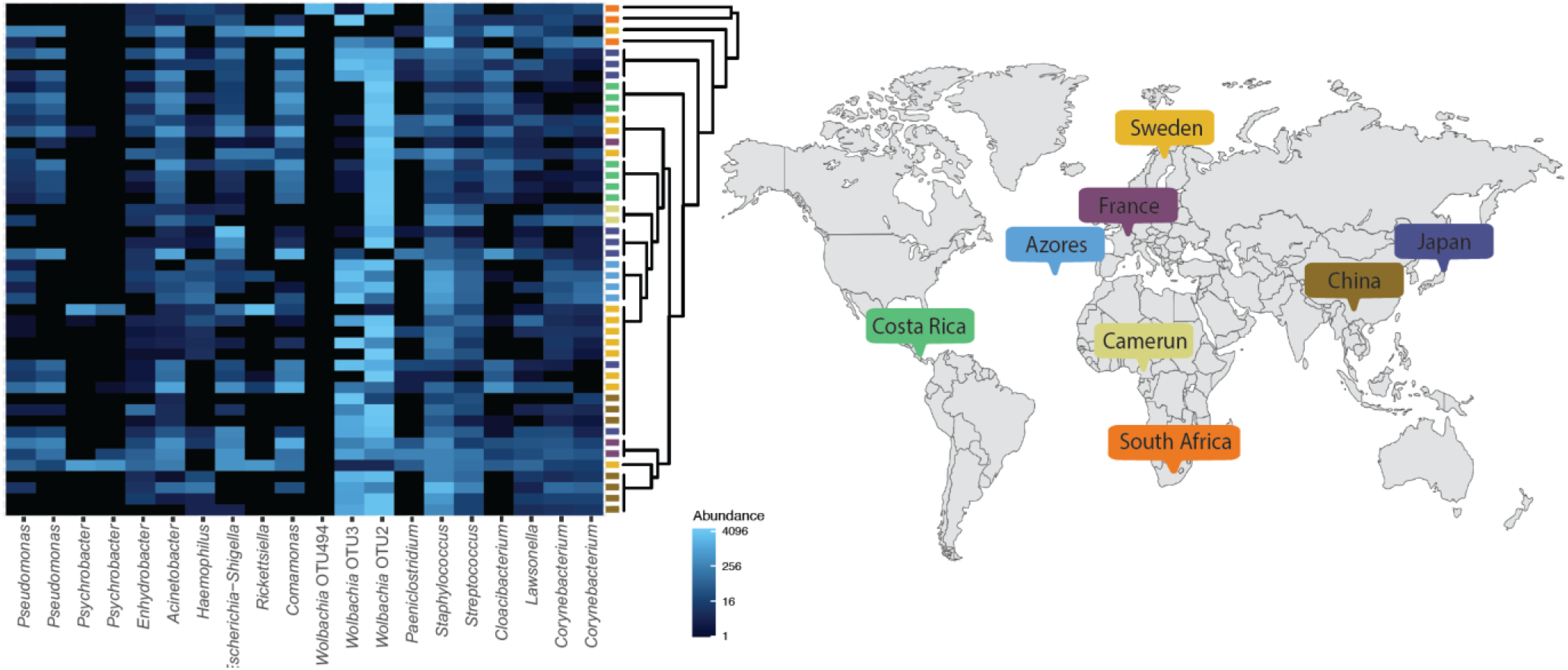
Compositional heat map produced for the 20 most abundant OTUs. The samples are ordered to reflect the phylogenetic relationships among analyzed *M. eurysternus* samples (complete COI phylogeny available in Supplementary figure 1), and color coded according to their geographical origin.

### Metagenomic assemblies and genomes characterization

Assembly of the *M. eurysternus* metagenomic data (sample GA72) contained two different *Wolbachia* strains. One strain (designated as *w*Meur1) was assembled into a single 732,857 bp long contig and closed into a complete circular genome using aTRAM extension to 733,850 bp. An identical sequence was obtained using PCR with specific primers. The second strain, *w*Meur2, was fragmented into 189 contigs. Screening of the 36 chewing lice metagenomic data available in SRA database revealed additional three strains of *Wolbachia*, designated hereafter as *w*Alce, wPaur, and *w*Mmer from *Alcedoecus* sp., *Penenirmus auritus*, and *Meromenopon meropis*, respectively. These genomes could only be assembled as drafts, composed of 9 (*w*Mmer) to 386 (*w*Alce) contigs. The BUSCO assessments indicate that these fragmented genomes are complete or almost complete (SupplementaryTable2). When assessed against Rickettsiales database, the average completeness was 95% (85.7%-98.7%). The assessment against Proteobacteria, performed to provide direct comparison with Lefoulon et al. (10), produced considerably lower average of 78.9% (68.5%-83.5%) corresponding to the results of Lefoulon et al. (10). For the *w*Alce strain, Busco predicted a higher degree of possible duplications, indicating that this assembly may not contain a single *Wolbachia* strain but could rather be a mixture of two closely related strains. *w*Meur1 and *w*Mmer display unusual, derived features. Particularly, *w*Meur1 is only 733,850 bp long with GC content of 28%. Based on RAST annotation it contains 592 protein coding genes, 3 rRNA genes, and 35 tRNAs. *w*Mmer genome is considerably longer (1,005,754 bp when concatenated) but with similarly low GC content (28.6%). Both genomes form long branches in phylogenetic trees (Figure 2, supplementary figures 3,4) and possess characteristics typical for obligate symbionts. Beside the low GC content, they do lack transposase sequences, phage-related sequences, and mobile elements. Ankyrin repeats are not present in *w*Meur1 and only one instance was detected in *w*Mmer (Table 1). The genomes of the remaining three strains resembled the previously reported genomes of supergroup F, containing transposase sequences, phage-related sequences, mobile elements, and ankyrin repeats (Table 1).

**Figure 2:**
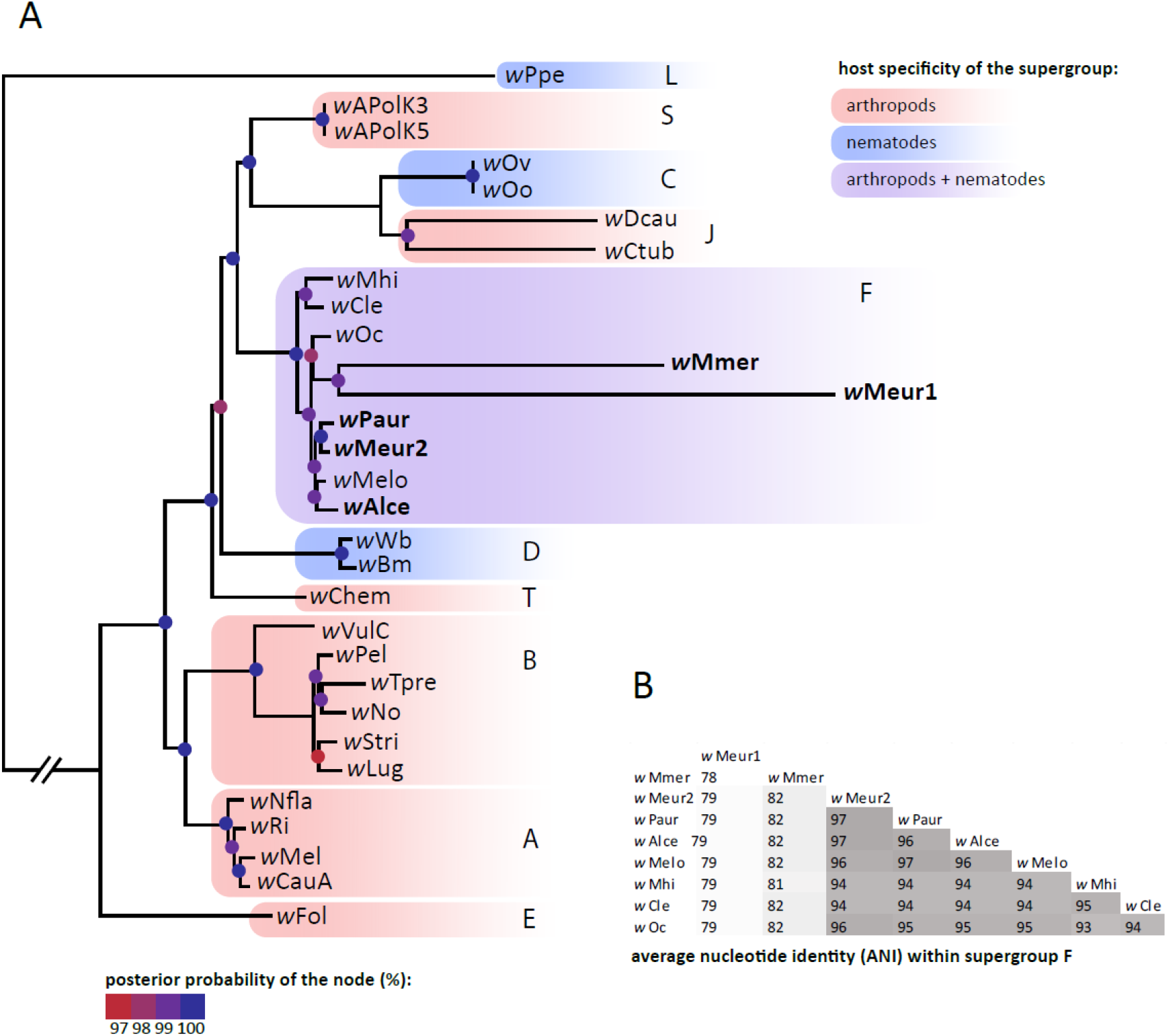
A - Phylogenetic relationships within the supergroup F derived for the available genomes (“multigene matrix” analyzed by Phylobayes). The new strains printed in bold. Posterior probabilities of the nodes are indicated by the colored dots. Super Supergroups are designated by the capital letters at the branches or clusters. B – Average nucleotide identity among the supergroup F genomes.

Orthofinder placed most of the protein coding genes from the five new strains into orthogroups shared with the other included strains from supergroup F. Overlap comparison between the genomes showed a high proportion of genes shared by all or most strains (Figure 3; Supplementary data 2). It also revealed that *w*Meur1 and *w*Mmer do not share any unique genes in exclusion of other genomes, despite their close phylogenetic relationship. On the other hand, while sharing a high proportion of genes, the strains displayed very limited degree of synteny (Supplementary figure 5). For example, Mauve analysis of closely related *w*Meur1 and *w*Mmer produced 159 local collinear blocks, the longest spanning 18 kb.

**Figure 3:**
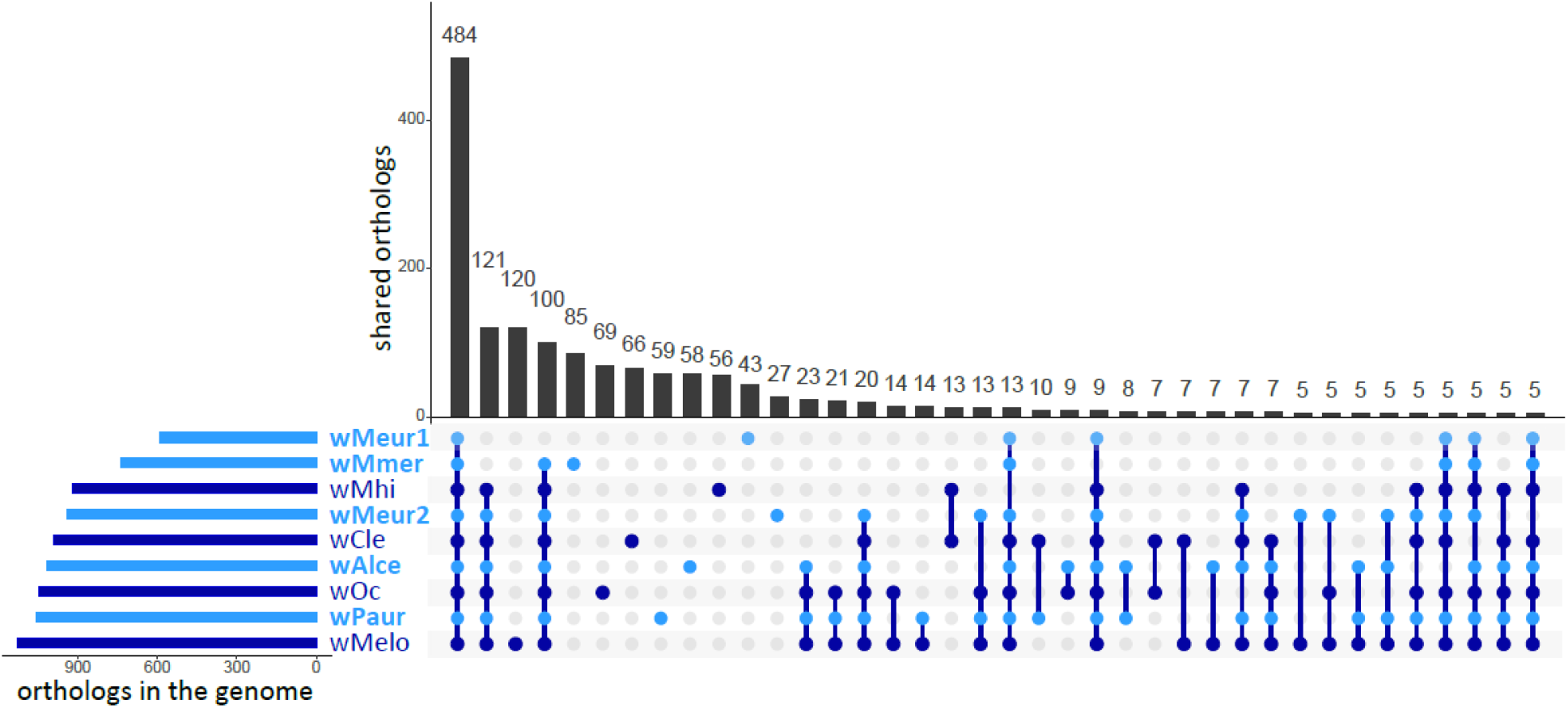
Orthogroups shared by the supergroup F Wolbachia genomes. Data for the new strains from chewing lice are printed in light blue.

### Horizontal transfer of genes for pantothenate synthesis

Orthofinder also identified a set of genes, each unique for a single strain, mostly annotated as hypothetical proteins (Supplementary table 4). A particularly interesting case is presented by three pantothenate synthesis related genes (panB, panG and panC) found exclusively in *w*Meur1. The only *Wolbachia* homologues found by diamond blastx were genes from *w*ClefT (NZ_CP051156), a strain infecting the cat flea *Ctenocephalides felis* (8). The other closest relatives originated from phylogenetically distant bacteria, including several symbiotic forms. In the phylogenetic analysis of the closest homologues retrieved by blast from NCBI, all three *w*Meur1 pantothenate genes clustered as sister taxa to their *w*CfeT homologues (Supplementary figure 6).

In *w*Meur1, the three pantothenate genes were the only instances of apparent horizontal transfers. In the other four chewing lice strains of *Wolbachia*, the search for HGT did not reveal horizontally transferred genes with recognized metabolic function. Majority of the genes provided blast hits from other *Wolbachia* strains, while the few instances of the non-*Wolbachia* origin included mostly ankyrins, transposases, and hypothetical proteins (SupplementaryData3).

### Comparison of metabolic capacities

Reconstruction of metabolic capacities shows high consistency across supergroup F genomes (Supplementary table 5). The main differences are found in the *w*Meur1 genome and, to a lesser extent, in the *w*Mmer genome. Both genomes lost a high portion of recombination genes and ABC transports. The smaller *w*Meur1 also lost genes for two B vitamin pathways retained in other strains (pyridoxine and folate) but acquired three genes for pantothenate synthesis by horizontal transfer (see above). All genomes show very limited capacity for amino acid synthesis. They only retain a near-complete pathway for lysine (leading probably to synthesis of peptidoglycan rather than lysine) and the glyA enzyme allowing for interconversion of serine and glycine.

### *Phylogenetic relationships of the* Wolbachia *symbionts*

All phylogenetic analyses of both matrices placed the newly described *Wolbachia* strains invariantly within the supergroup F (Figure 2, Supplementary figures 3,4). The two most derived strains *w*Meur1 and *w*Mmer formed extremely long branches within the supergroup, comparable only to those of the nematode-associated mutualists from the supergroups J. In ML analysis of the “*multigene*” matrix these two long branches clustered at the base of the supergroup F (Supplementary figure 4), while in Phylobayes analysis they were nested among the other supergroup F taxa (Figure 2). The remaining three strains, *w*Meur2, *w*Alce, *w*Paur, were placed on considerably shorter branches, comparable to the rest of *Wolbachia* included in the analysis.

## Discussion

### New highly derived members of supergroup F

The new *Wolbachia* supergroup F strains described here from four species of chewing lice represent two remarkably different types of symbiont genomes. Three of them (*w*Meur2, *w*Paur, *w*Alce) resemble the other *Wolbachia* strains of supergroup F in their genomic characteristics (size above 1 Mb, average GC content between 35% and 37%, presence of phage-related sequences, mobile elements, transposases, ankyrin repeats, etc.). This close similarity is also reflected in the short branches they form in the phylogenetic trees (Figure 2, Supplementary figures 3,4). However, two additional strains (*w*Meur1, *w*Mmer) show very specific and derived traits, unique within the context of the whole *Wolbachia* diversity. Particularly, *w*Meur1 represents the first highly reduced, insect-associated *Wolbachia* strain with characteristics typical for obligate mutualists known from other bacterial groups (20, 57, 58). It also provides another candidate example of transition towards mutualism by horizontal gene transfer (20, 59). This modifies the view that the arthropod associated strains of *Wolbachia* generally possess larger genomes, richer with transposable elements, prophage-related genes or repeat-motif proteins than their nematode-associated relatives (4, 60). With its 733,850 bp length *w*Meur1 is currently the smallest known genome among *Wolbachia*, almost 100 kb shorter than its nematode-associated “predecessors” *w*Ctub and *w*Dcau (10). The second derived strain, *w*Mmer, possesses a genome larger than *w*Meur1, but shows similar signs of strong degeneration: both strains lack phage-related sequences, mobile elements, transposases and ankyrin repeats (with exception of one ankyrin repeat found in *w*Mmer).

### Phylogenetic relationships

The highly derived nature of *w*Meur1 and *w*Mmer genomes is also apparent from the low ANI values, when compared to their close relatives, and the long branches they form in phylogenetic trees (Figure 2). In principle, such long branches may distort phylogenetic analyses. This phenomenon is particularly dangerous when analyzed taxa differ considerably in their nucleotide composition, e.g., when symbiotic genomes with extremely low GC content are included. In our study, the placement of these two symbionts within supergroup F is supported by several indications. First, this placement is not likely to be affected by long branches since no other members of supergroup F possess long branches which would attract the *w*Meur1+*w*Mmer pair. Second, the position of the two genomes as sister taxa has been retrieved with high support from all analyses, including the phylobayes analysis which is particularly resistant to this type of artifact (52).

Phylogenetic relationships within supergroup F suggest that at least the short-branched *Wolbachia* strains were acquired independently by their chewing lice hosts. For example, the two strains from philopterid lice, *w*Alce and *w*Paur, do not cluster as sister taxa in any of the analyses. While the tree resolution is rather poor and does not provide clear evidence, horizontal transfers within the F supergroup have been deduced previously, e.g., between isopods and termites (13). This apparent lack of a coevolutionary signal is also concurrent with the absence of *Wolbachia* in all other screened SRA data for chewing lice (discussed below), and with the presence of a phylogenetically distant gammaproteobacterial symbiont in *Columbicola wolffhuegeli* (26, 61).

### Distribution of the Wolbachia in chewing lice species

In our study, we found *Wolbachia* contigs in five out of the 36 assemblies of the SRA datasets. This is in sharp contrast with the results reported by Kyei-Poku et al. (14). In their screening, focused on *Wolbachia* in lice, these authors showed the presence of these bacteria in all 19 tested species (interestingly, none of them from supergroup F, all falling into A and B). However, their approach was based on PCR amplification of selected genes using specific primers. It is likely that this method can detect *Wolbachia* even if present in extremely low numbers. In contrast, the WGS-based approach will only produce data for the dominant bacteria in the microbiome. This is also reflected in the amplicon analyses, which revealed considerably more bacterial taxa in each individual *M. eurysternus* sample (Figure 1). This ambiguity raises the question, which of the *Wolbachia* previously detected in lice are parasites (known to be broadly distributed across arthropod taxa) and which may possibly represent the comparatively rarer instance of obligate mutualist.

### *Highly reduced* w*Meur1: transition to obligate symbiosis by HGT of pantothenate synthesis genes?*

The genomic comparisons of the new strains are entirely consistent with their phylogenetic patterns. While the short-branched strains are closely similar in their genome characteristics, the two most derived genomes (*w*Meur1 and *w*Mmer) differ in many aspects. They both underwent considerable deterioration of the recombination and transport systems and unlike the other strains, they lack the elements related to the genomic dynamics and fluidity (phages, transposases, mobile elements). Such evolutionary trends accompanying the reduction in genome size are well known from many obligate insect symbionts from gammaproteobacteria (62) but are much less common in *Wolbachia*. For example, several *Wolbachia* strains have been suggested to establish mutualistic relationships with their hosts after acquiring complete biotin operon (8, 59). Within insect hosts, these systems include *w*Cle in the bedbug *Cimex lectularius, w*CfeT in the flea *Ctenocephalides felis* (8), or two *Wolbachia* strains from *Nomada* bees (63). In the latter, *Wolbachia* phylogeny even suggests co-divergence across several host species, typical for mutualistic obligate symbionts. However, in all these insects, *Wolbachia* symbionts retain genomes which exceed 1Mb and contain many mobile elements. As pointed out by Driscoll et al. (8), *w*Cle and *w*CfeT even possess extremely high numbers of pseudogenes (see table1 for *w*Cle). The *w*Meur1 strain presented here differs from these insect symbionts by a dramatic reduction and “cleansing” of its genome. It is not only the smallest known genome among *Wolbachia* but also the first insect *Wolbachia* with genomic characteristics typical for obligate mutualists.

These features bring into question the role of the *w*Meur1 strain in its host. Unlike the above examples of presumably mutualistic *Wolbachia*, the *w*Meur1 strain does not possess genes required for biotin synthesis. Of the other vitamin B pathways, often considered to be of potential importance in nutritional symbionts, it only retains the production of riboflavin, a pathway conserved across many *Wolbachia* strains (64). However, a striking metabolic difference between *w*Meur1 and all other supergroup F strains is the presence of three genes required for synthesis of pantothenate, most likely acquired by horizontal gene transfer (HGT). The blast-based HGT analysis retrieved the most closely similar homologue from *w*CfeT, a *Wolbachia* strain from cat flea *Ctenocephalides felis*, while the other retrieved homologues belonged to other, often phylogenetically distant bacterial groups (Supplementary data 3). In the phylogenetic analysis, all three genes from *w*Meur1 and *w*CfeT formed closely related sister taxa (Supplementary figure 6). Remarkably, the same triad of genes is also present in the genome of a *Sodalis*-related symbiont from another chewing louse, *C. wolffhuegeli* (Figure 4; NCBI bioproject PRJNA692390). When characterizing metabolic capacity of this bacterium, Alickovic et al. (26) mainly addressed the issue of keratin digestion and concluded that no clear metabolic role can be deduced from the symbiont’s genome content. Similarly, in our new strains, we failed to detect production of enzymes with keratinase activity, and we observed almost complete loss of capacity for amino acid synthesis (Supplementary table 5). However, the presence of the three pantothenate related genes in *C. wolffhuegeli* symbiont and their HGT acquisition by *w*Meur1 suggests that production of this vitamin might be at least part of the metabolic function in these obligate mutualists.

**Figure 4:**
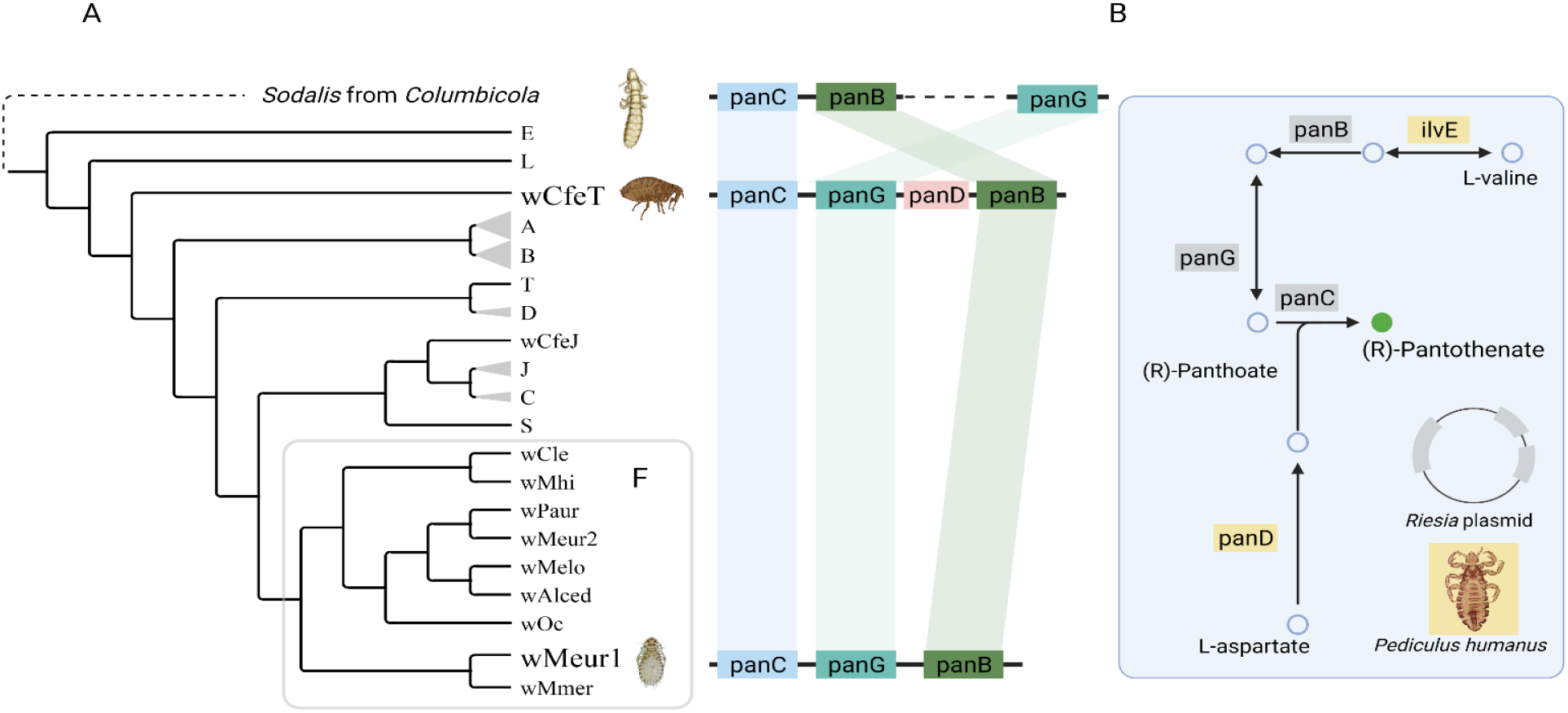
Distribution of the pantothenate synthesis related genes in the *Wolbachia* genomes. A – The genes mapped on a schematic phylogeny tree. B – overview of the pantothene synthesis pathway in *Pediculus humanus* and its symbiont *Riesia pediculicola* (reconstruction based on KEGG database).

While the triad panB, panG, and panC forms a core of pantothenate synthesis, full functionality of the pathway requires two additional genes, ilvE and panD (Figure 4). These genes are missing in the *w*Meur1 genome but are very likely present in the genome of the host. The phylogenetically closest system with fully characterized genomic capacities is the symbiosis between human louse *Pediculus humanus* and its symbiont *Riesia pediculicola* (65). In this blood feeding insect, *Riesia* provides the same three genes (panB, panG and panC located on plasmid) while ilvE and panD are present in the host genome (in Figure 4 shown on simplified reconstruction based on the KEGG database). Our blast screening of the *M. eurysternus* assembly shows that these genes are also present in the *w*Meur1 host and suggests the complete pantothenate pathway is therefore functional.

According to the hypothetical scenario introduced by Lo et al. (66), symbiogenesis involves two bouts of dramatic genomic changes. The first occurs during the transition from free-living bacterium to a facultative symbiont and the second one with transition to an obligate symbiont. During these processes, bacterial genomes first undergo a dramatic decrease of coding density due to large-scale inactivation of the genes. This step is followed by removal of the inactivated genes and restoration of the coding density. Our set of new *Wolbachia* strains fit well into this scenario. As shown in Figure 5 most of the supergroup F genomes display relatively high coding density between 78% and 84%. These genomes carry transposases and mobile elements, and with exception of *w*Cle, also a relatively rich repertoire of recombination/repair genes (Supplementary data 4). In contrast, the *w*Mmer genome shows a considerable drop in coding density to 65%. While its size is comparable to the former strains, it contains a significantly lower gene number, indicating large-scale deactivation. Finally, the *w*Meur1 genome restores the coding density to 76%, apparently due to removal of its deactivated regions. Its position in this evolutionary spectrum, loss of most of the recombination/repair systems, and presumed metabolic role in pantothenate provision provide strong evidence that *w*Meur1 is the first known insect-associated *Wolbachia* strain which completed the transition to obligate mutualist.

**Figure 5:**
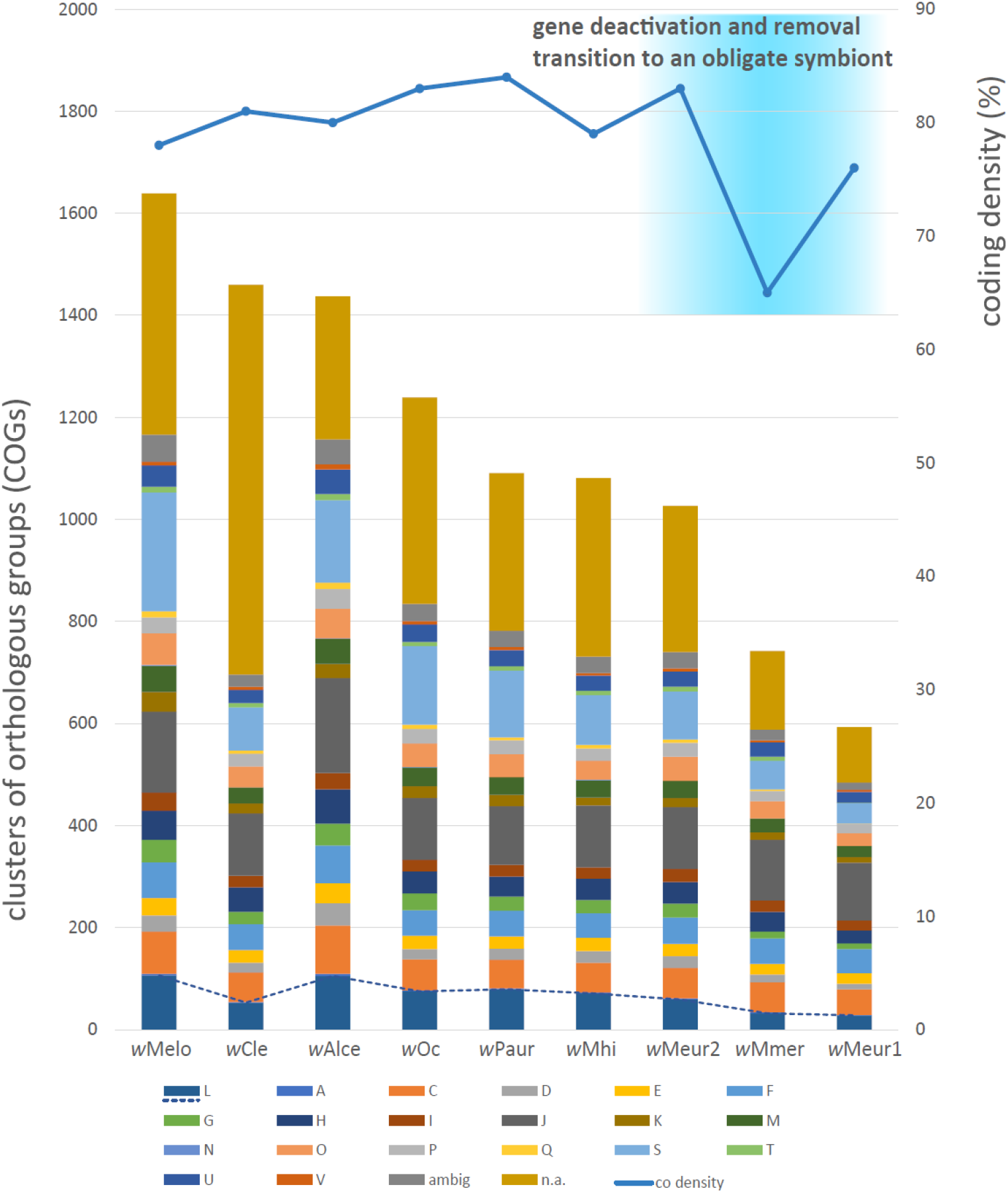
Comparison of the supergrup F Wolbachia genomes in respect to coding density and numbers of genes in different COGs. A - RNA processing and modification, C - Energy production and conversion, D - Cell cycle control and mitosis, E - Amino Acid metabolis and transport, F - Nucleotide metabolism and transport, G - Carbohydrate metabolism and transport, H - Coenzyme metabolism, I - Lipid metabolism, J - Translation, K - Transcription, L - Replication, recombination and repair, M - Cell wall/membrane/envelop biogenesis, N - Cell motility, O - Post-translational modification, protein turnover, chaperone functions, P - Inorganic ion transport and metabolism, Q - Secondary Structure, T - Signal Transduction, U - Intracellular trafficing and secretion, V - Defense mechanisms, S - Function Unknown, ambig - assigned to more than one category, n.a. – not assigned, co density - curve showing coding density. Dashed line highlights differences in the L category related to replication, recombination and repair.

## Acknowledgments

This work was supported by the Grant Agency of the Czech Republic (grant 20-07674S to V.H.). We would like to acknowledge the sequencing services of the DNA Services of Roy J. Carver Biotechnology Center, University of Illinois at Urbana-Champaign, Illinois, USA. Access to computing and storage facilities owned by parties and projects contributing to the National Grid Infrastructure MetaCentrum provided under the programme “Projects of Large Research, Development, and Innovations Infrastructures” (CESNET LM2015042), is greatly appreciated. We thank Joel J. Brown for his language corrections, and Daniel R. Gustafsson (Guangdong Academy of Sciences) for providing us samples of *M. eurysternus* from China, France, Japan and Sweden for amplicon analysis.

