## Supplementary figure 1 for "Extremely reduced supergroup F *Wolbachia*: transition to obligate insect symbionts"

**Supplementary figure 1:** host phylogeny - relationships of *Menacathus eurysternus* samples. The sample used in this study for the metagenomic assembly printed in bold blue.

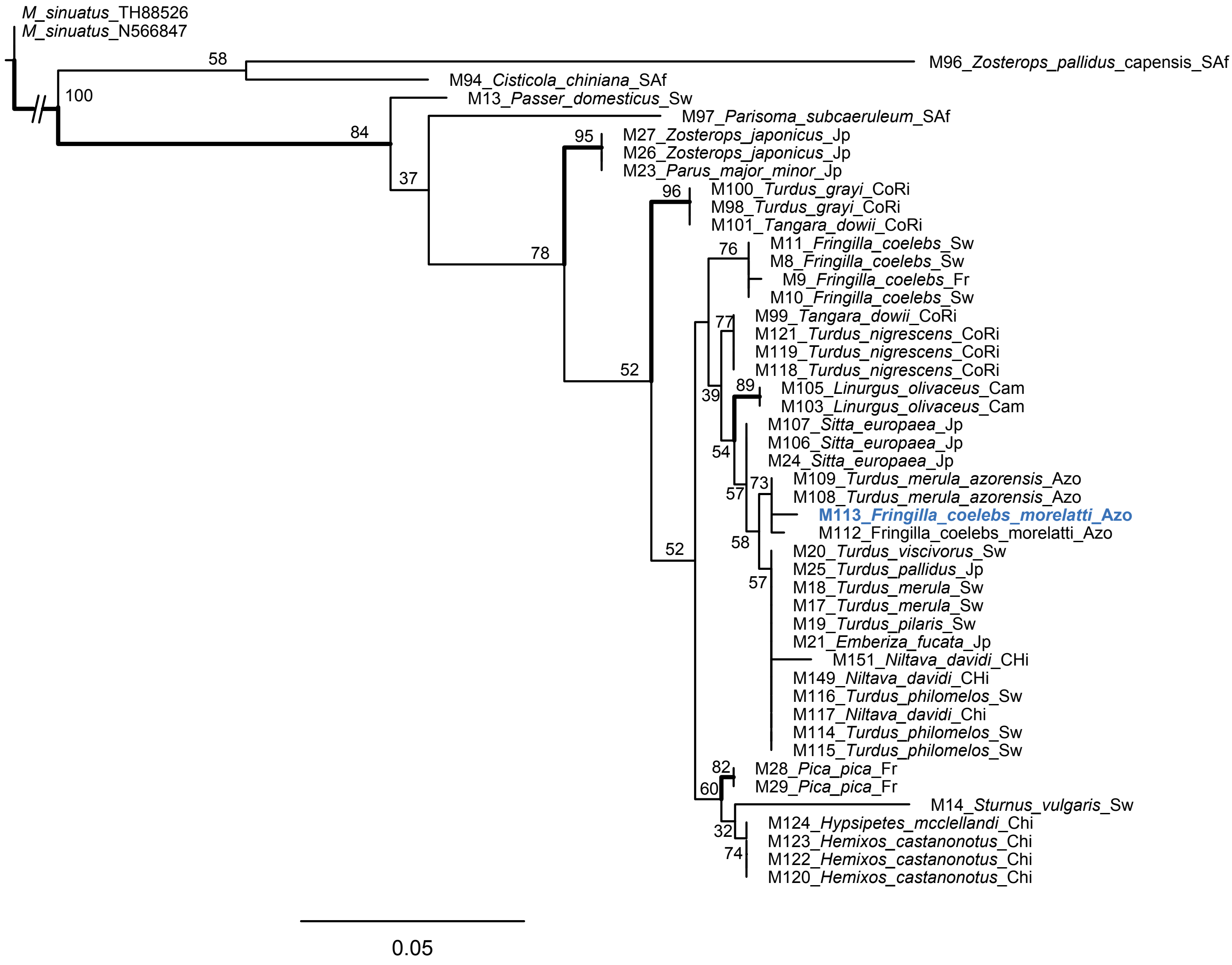
