## Supplementary figure 2 for "Extremely reduced supergroup F *Wolbachia*: transition to obligate insect symbionts"

**Supplementary figure 2:** Compositional heat map for M. eurysternus microbiomes based on the strictly decontaminated 16S rRNA dataset (see Materials and Methods) rarefied at 1000 (A) and 2000 reads (B). The sample order reflects Figure 1 in the main manuscript. Additional information on the samples are found in Supplementary Data1.

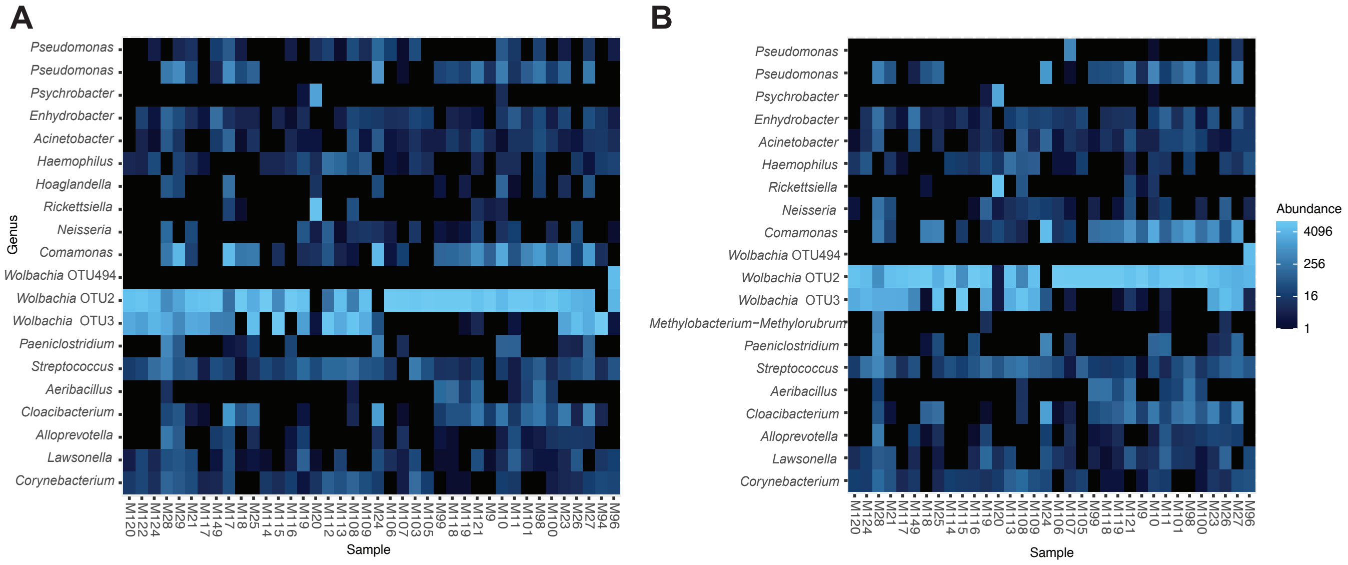
