## Supplementary figure 3 for "Extremely reduced supergroup F *Wolbachia*: transition to obligate insect symbionts"

**Supplementary figure 3:** Phylogenetic tree derived from the two-gene matrix by IQ-TREE. The genomes assembled in this study printed in bold blue.

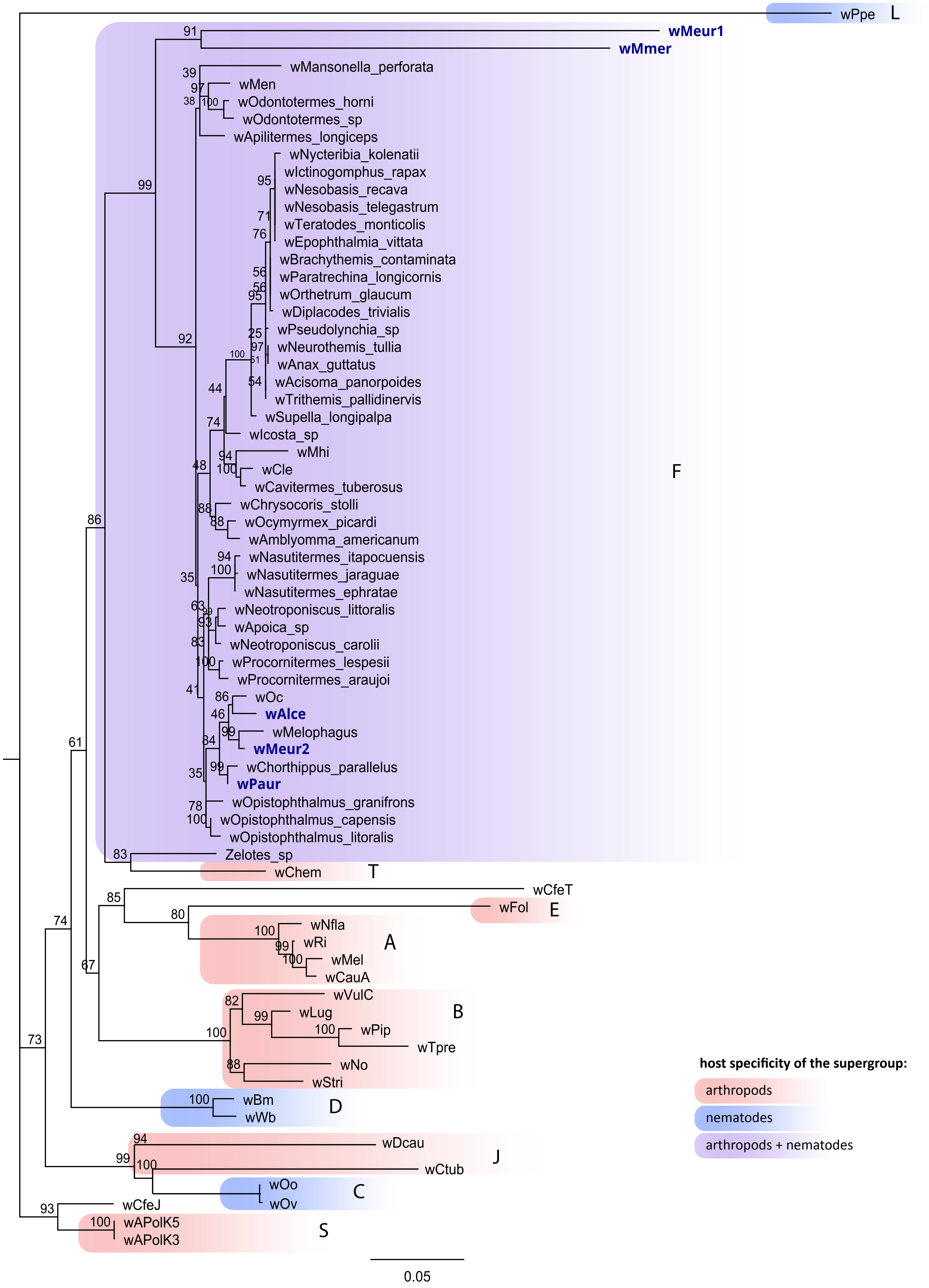
