## Supplementary figure 4 for "Extremely reduced supergroup F *Wolbachia*: transition to obligate insect symbionts"

**Supplementary figure 4:** Phylogenetic trees derived from the multigene matrix by ML. The genomes assembled in this study printed in bold blue.

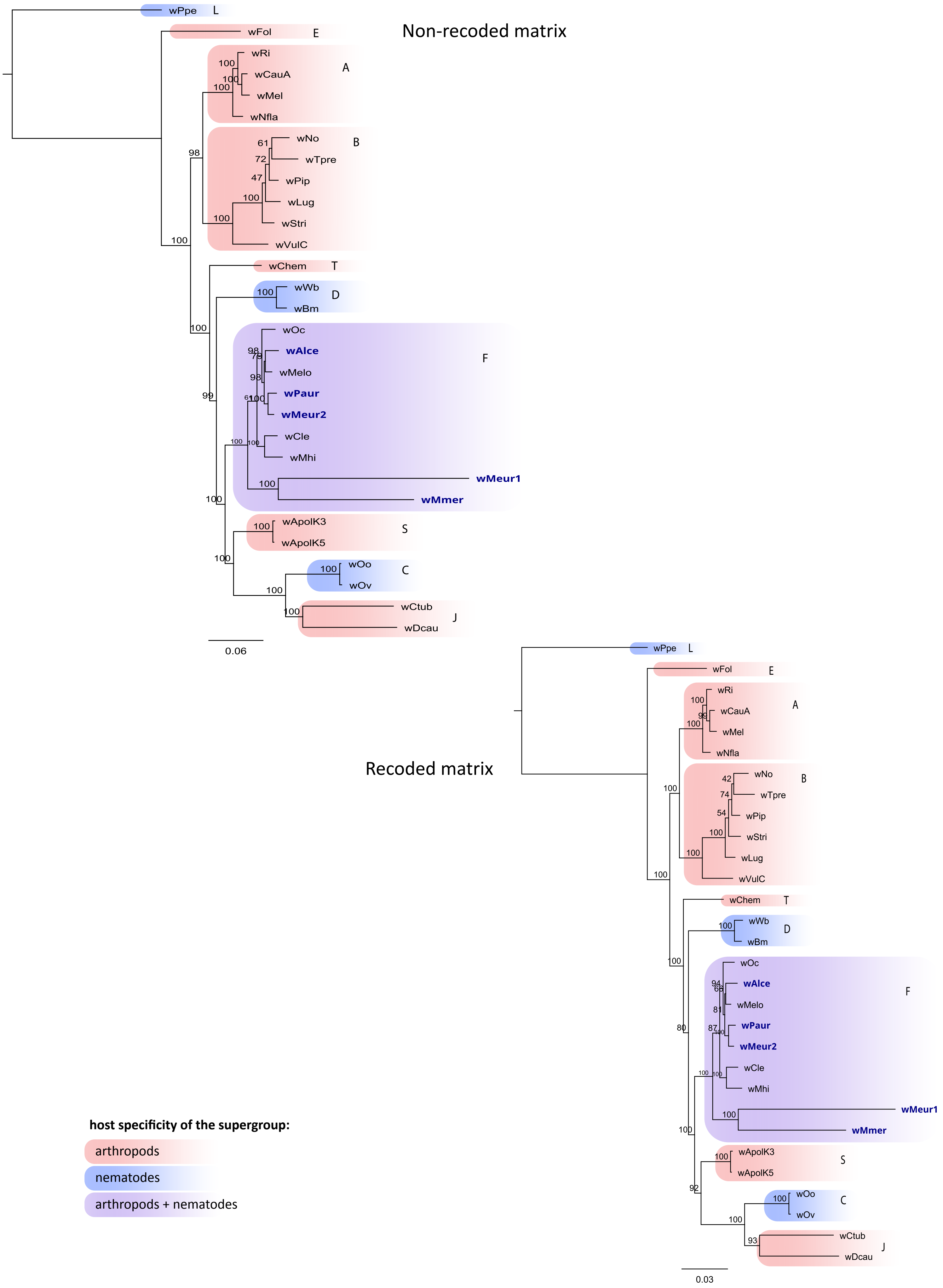
