## Supplementary figures and images for "Extremely reduced supergroup F *Wolbachia*: transition to obligate insect symbionts"

### Supplementary figure 5

**Supplementary figure 5:** Mauve synteny analysis.

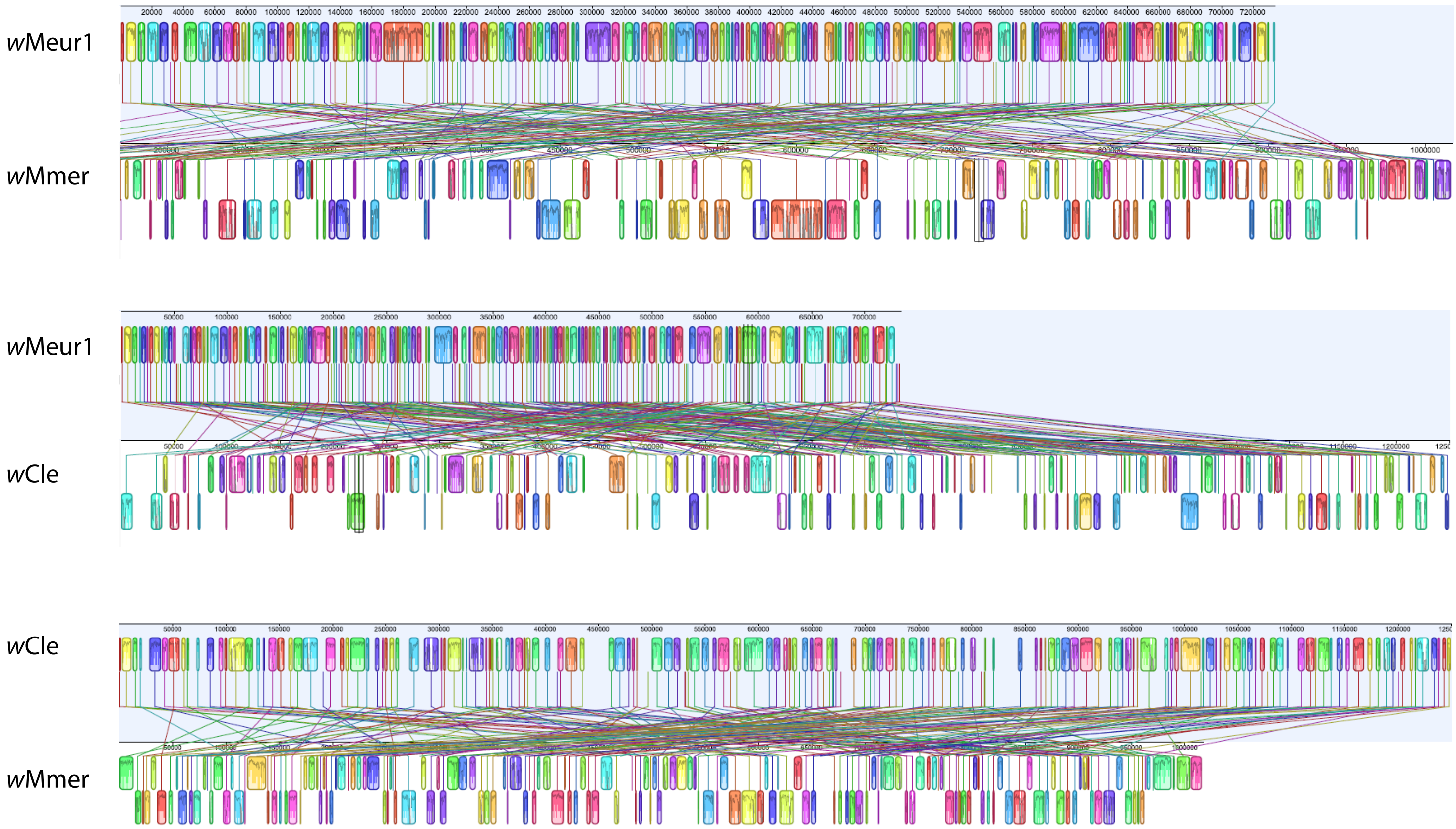
