## Supplementary figure 6 for "Extremely reduced supergroup F *Wolbachia*: transition to obligate insect symbionts"

**Supplementary figure 6A**: Phylogenetic position of pantothenate synthesis related genes; Pantoate--beta-alanine ligase panC (EC 6.3.2.1)

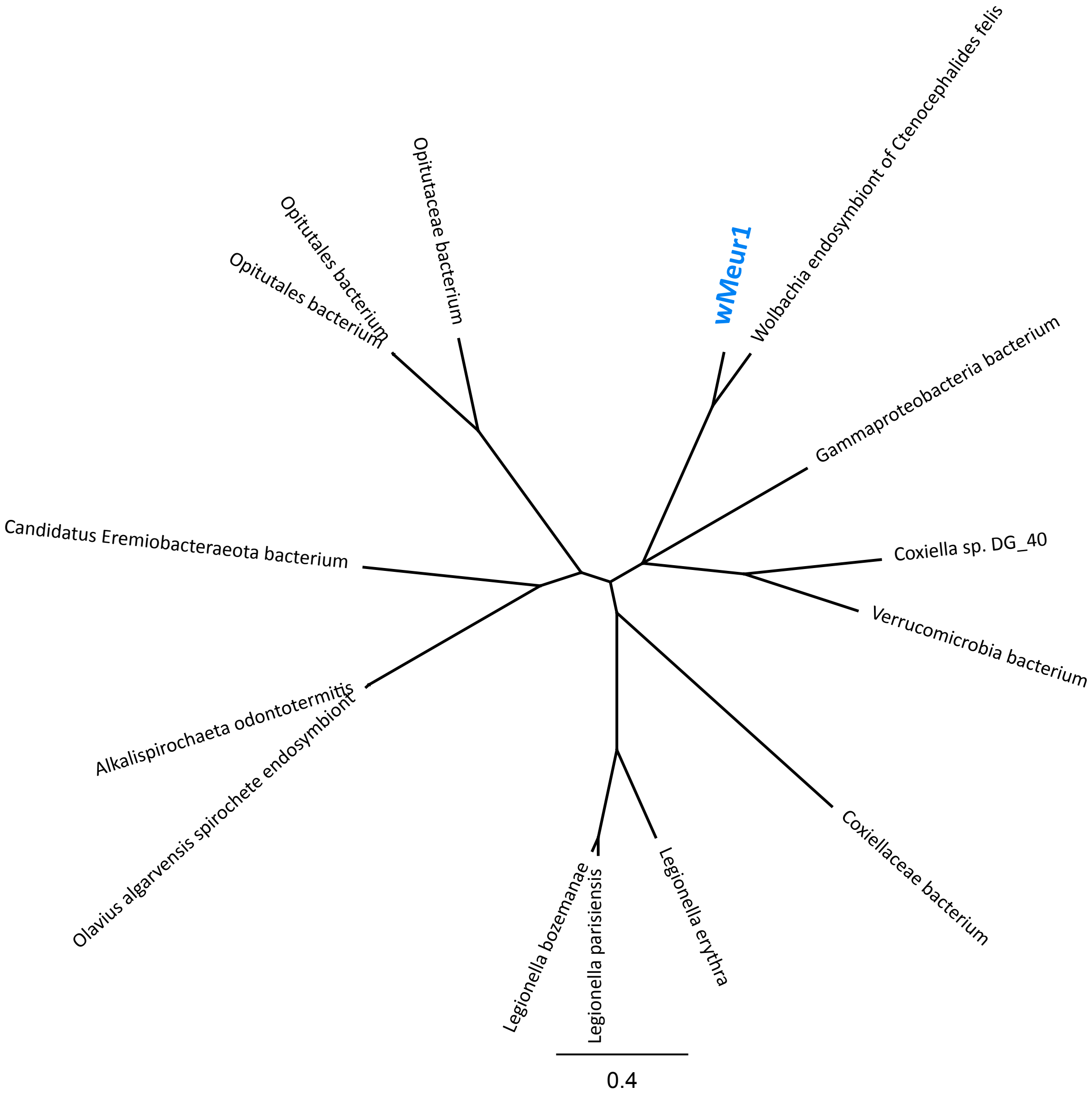


| **Organism** | **Name** |
|---|---|
| Olavius algarvensis spirochete endosymbiont | WP_124032323 |
| Alkalispirochaeta odontotermitis | WP_037564257 |
| Candidatus Eremiobacteraeota bacterium | MBN9417370 |
| Opitutaceae bacterium | MAM92216 |
| Opitutales bacterium | MBT6769675 |
| Opitutales bacterium | MBT3482533 |
| Coxiellaceae bacterium | MBV53023 |
| Legionella erythra | WP_058527648 |
| Legionella parisiensis | WP_058516236 |
| Legionella bozemanae | WP_058458018 |
| Wolbachia endosymbiont of Ctenocephalides felis wCfeT | WP_168464627 |
| Gammaproteobacteria bacterium | MBI2790727 |
| Coxiella sp. DG_40 | KPJ67214 |
| Verrucomicrobia bacterium | PWU05281 |

**Supplementary figure 6B**: Phylogenetic position of pantothenate synthesis related genes; Ketopantoate reductase PanG (EC 1.1.1.169)

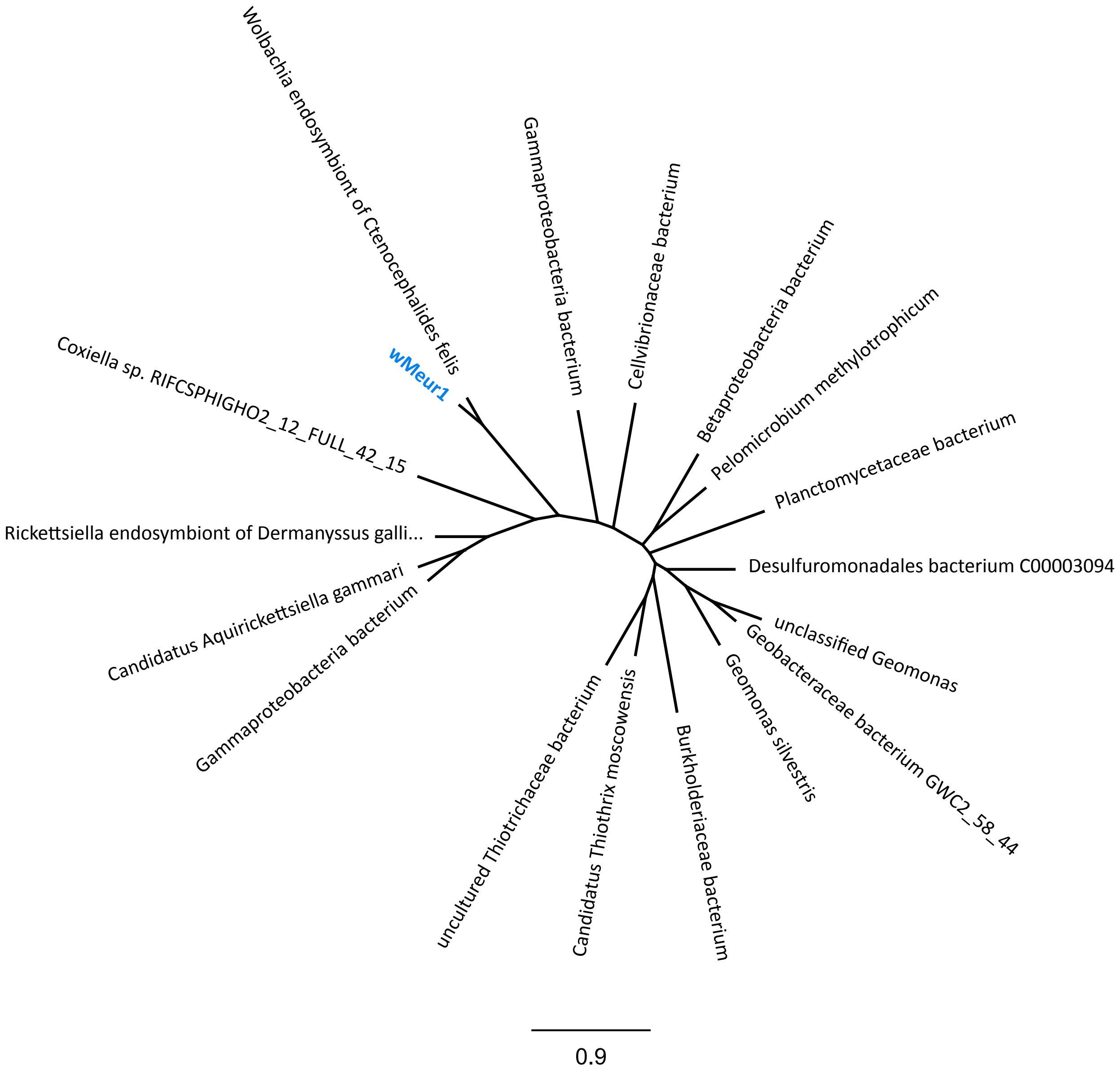


| Name | Accession |
|---|---|
| Gammaproteobacteria bacterium | TLY47062 |
| Candidatus Aquirickettsiella gammari | RDH40168 |
| Rickettsiella endosymbiont of Dermanyssus gallinae | WP_218814024 |
| Wolbachia endosymbiont of Ctenocephalides felis wCfeT | WP_218938889 |
| Gammaproteobacteria bacterium | HBA33227 |
| Burkholderiaceae bacterium | MBV8634101 |
| Cellvibrionaceae bacterium | MAZ88707 |
| Planctomycetaceae bacterium | HID78016 |
| Betaproteobacteria bacterium | MBI1965041 |
| Pelomicrobium methylotrophicum | WP_147800823 |
| Desulfuromonadales bacterium C00003094 | OEU60518 |
| uncultured Thiotrichaceae bacterium | CAA6822986 |
| Candidatus Thiothrix moscowensis | MBJ6612093 |
| Geomonas silvestris | WP_183355568 |
| Geobacteraceae bacterium GWC2_58_44 | OGU05969 |
| unclassified Geomonas | WP_217289478 |
| Coxiella sp. RIFCSPHIGHO2_12_FULL_42_15 | OGO93865 |

**Supplementary figure 6C:** Phylogenetic position of pantothenate synthesis related genes; 3-methyl-2-oxobutanoate hydroxymethyltransferase pan B (EC 2.1.2.11)

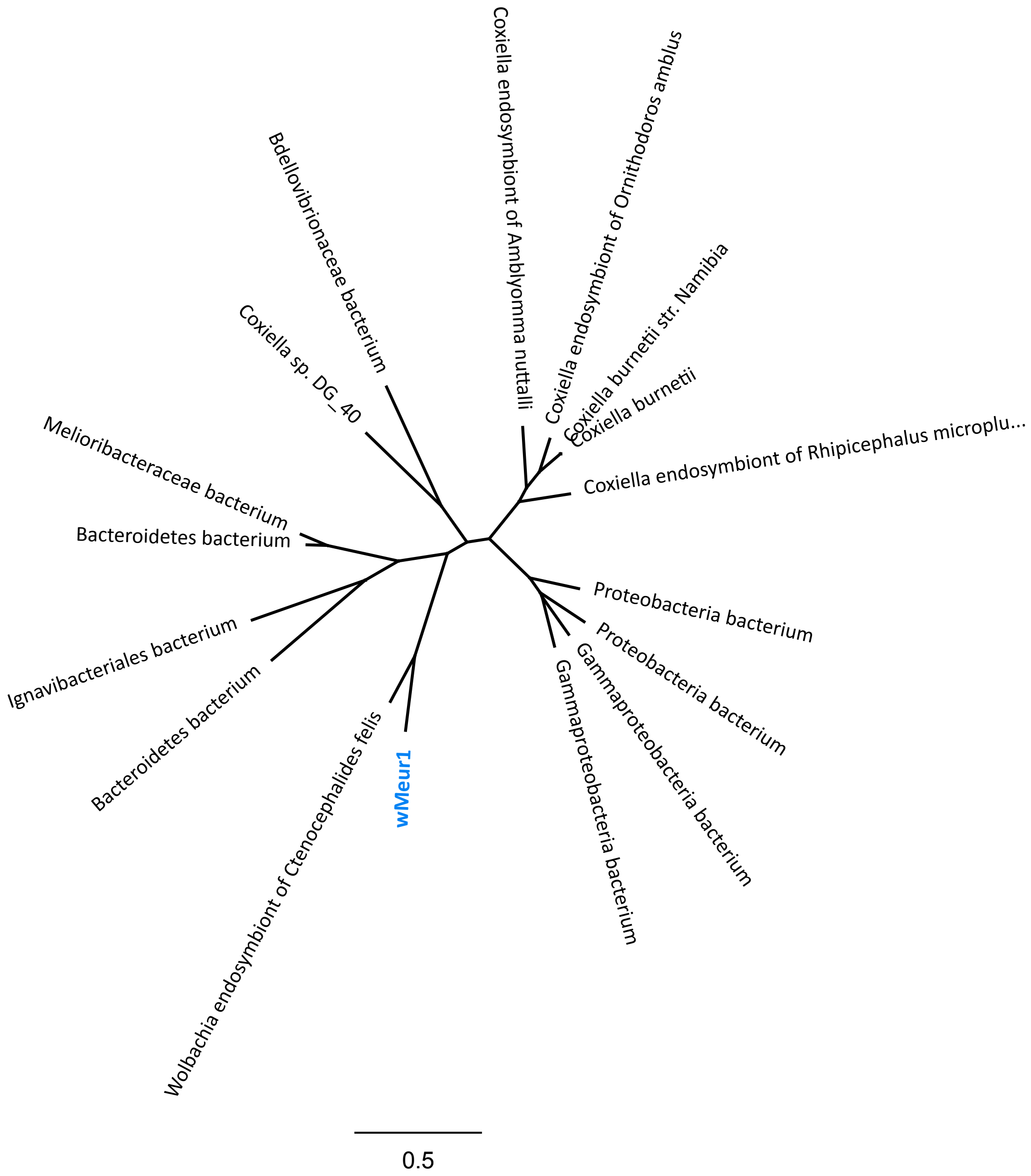


| Name | Accession |
|---|---|
| Bacteroidetes bacterium | MBU0473530 |
| Ignavibacteriales bacterium | MBN2571901 |
| Melioribacteraceae bacterium | KAB2839543 |
| Bacteroidetes bacterium | MBU2494347 |
| Wolbachia endosymbiont of Ctenocephalides felis wCfeT | WP_168464630 |
| Coxiella sp. DG_40 | KPJ67213 |
| Proteobacteria bacterium | MBS0288437 |
| Proteobacteria bacterium | MBS0287918 |
| Gammaproteobacteria bacterium | MBI2790726 |
| Gammaproteobacteria bacterium | MBN9286951 |
| Coxiella endosymbiont of Rhipicephalus microplus | WP_102157157 |
| Coxiella endosymbiont of Amblyomma nuttalli | WP_211923626 |
| Coxiella endosymbiont of Ornithodoros amblus | MBW5802677 |
| Coxiella burnetii str. Namibia | AIT63869 |
| Coxiella burnetii | WP_080744740 |
| Bdellovibrionaceae bacterium | MBT4762628 |
