## Supplementary table 1 for "Extremely reduced supergroup F *Wolbachia*: transition to obligate insect symbionts"

**Supplementary table 1**: SRA samples assembled and screened for *Wolbachia*. Highlighted by blue = positive screening resulting in new strains described in this study. Highlighted by grey = weak positive screening, not included into this study.

| SRA sample | Source | Sub-order/Family | Vertebrate host | strain |
|---|---|---|---|---|
| SRR5308125 | *Heterodoxus spiniger* | Amblycera/Boopiidae | *Canis lupus* | |
| SRR5308130 | *Macrogyropus costalimai* | Amblycera/Gyropidae | *Cuniculus paca* | |
| SRR5308127 | *Laemobothrion tinnunculi* | Amblycera/Laemobothriidae | *Falco longipennis* | |
| SRR5088470 | *Osborniella crotophagae* | Amblycera/Menoponidae | *Crotophaga ani* | |
| SRR5308132 | *Myrsidea* sp. | Amblycera/Menoponidae | *Myiothlypis luteoviridis* | |
| SRR8145978 | *Menopon gallinae* | Amblycera/Menoponidae | *Gallus gallus* | |
| SRR8334265 | *Meromenopon meropis* | Amblycera/Menoponidae | *Merops apiaster* | *w* Mmer |
| SRR5308140 | *Ricinus* sp. | Amblycera/Ricinidae | *Myiothlypis luteoviridis* | |
| SRR5308146 | *Cummingsia maculata* | Amblycera/Trimenoponidae | *Lestoros inca* | |
| SRR5088466 | *Bothriometopus macrocnemis* | Ischnocera/Philopteridae | *Chauna torquata* | |
| SRR5308110 | *Alcedoecus* sp. | Ischnocera/Philopteridae | *Halcyon badia* | *w* Alce |
| SRR5308111 | *Anatoecus icterodes* | Ischnocera/Philopteridae | *Anas cyanoptera* | |
| SRR5308112 | *Brueelia antiqua* | Ischnocera/Philopteridae | *Catharus ustulatus* | |
| SRR5308113 | *Campanulotes compar* | Ischnocera/Philopteridae | *Columba livia* | |
| SRR5308114 | *Chelopistes texanus* | Ischnocera/Philopteridae | *Ortalis vetula* | |
| SRR5308115 | *Columbicola columbae* | Ischnocera/Philopteridae | *Columba livia* | |
| SRR5308116 | *Craspedonirmus immer* | Ischnocera/Philopteridae | *Gavia immer* | |
| SRR5308117 | *Docophoroides brevis* | Ischnocera/Philopteridae | *Diomedea exulans* | |
| SRR5308118 | *Falcolipeurus marginalis* | Ischnocera/Philopteridae | *Cathartes aura* | |
| SRR5308119 | *Fulicoffula longipila* | Ischnocera/Philopteridae | *Fulica americana* | |
| SRR5308120 | *Goniodes ortygis* | Ischnocera/Philopteridae | *Colinus virginianus* | |
| SRR5308124 | *Halipeurus diversus* | Ischnocera/Philopteridae | *Puffinus tenuirostris* | |
| SRR5308126 | *Ibidoecus bisignatus* | Ischnocera/Philopteridae | *Plegadis chihi* | |
| SRR5308131 | *Megaginus tataupensis* | Ischnocera/Philopteridae | *Crypturellus tataupa* | |
| SRR5308133 | *Osculotes curta* | Ischnocera/Philopteridae | *Opisthocomus hoazin* | |
| SRR5308134 | *Oxylipeurus chiniri* | Ischnocera/Philopteridae | *Ortalis vetula* | |
| SRR5308135 | *Pectinopygus varius* | Ischnocera/Philopteridae | *Phalacrocorax varius* | |
| SRR5308137 | *Penenirmus auritus* | Ischnocera/Philopteridae | *Sphyrapicus varius* | *w* Paur |
| SRR5308139 | *Quadraceps punctatus* | Ischnocera/Philopteridae | *Larus argentatus* | |
| SRR5308141 | *Saemundssonia lari* | Ischnocera/Philopteridae | *Larus novaehollandiae* | |
| SRR5308142 | *Strongylocotes lipogonus* | Ischnocera/Philopteridae | *Rhynchotus rufescens* | |
| SRR5308144 | *Trichophilopterus babakotophilus* | Ischnocera/Philopteridae | *Propithecus verreauxi* | |
| SRR5308145 | *Pessoaiella absita* | Ischnocera/Philopteridae | *Opisthocomus hoazin* | |
| SRR1821919 | *Geomydoecus ewingi* | Ischnocera/Trichodectidae | not provided | |
| SRR5308121 | *Geomydoecus aurei* | Ischnocera/Trichodectidae | *Thomomys bottae* | |
| SRR5308143 | *Stachiella larseni* | Ischnocera/Trichodectidae | *Mustela vison* | |
