## Supplementary table 2 for "Extremely reduced supergroup F *Wolbachia*: transition to obligate insect symbionts"

**Supplementary table 2**: Busco evaluation of the genome's completeness; high level of duplication in *w*Alce highlited by grey background.

**Rickettsia database**

wMeur1
C:85.7%[S:85.7%,D:0.0%],F:0.3%,M:14.0%,n:364
312 Complete BUSCOs (C)
312 Complete and single-copy BUSCOs (S)
0 Complete and duplicated BUSCOs (D)
1 Fragmented BUSCOs (F)
51 Missing BUSCOs (M)
364 Total BUSCO groups searched

wMmer
C:91.8%[S:91.8%,D:0.0%],F:0.3%,M:7.9%,n:364
334 Complete BUSCOs (C)
334 Complete and single-copy BUSCOs (S)
0 Complete and duplicated BUSCOs (D)
1 Fragmented BUSCOs (F)
29 Missing BUSCOs (M)
364 Total BUSCO groups searched

wMeur2
C:98.6%[S:98.6%,D:0.0%],F:0.3%,M:1.1%,n:364
359 Complete BUSCOs (C)
359 Complete and single-copy BUSCOs (S)
0 Complete and duplicated BUSCOs (D)
1 Fragmented BUSCOs (F)
4 Missing BUSCOs (M)
364 Total BUSCO groups searched

wPaur
C:97.0%[S:97.0%,D:0.0%],F:1.9%,M:1.1%,n:364
353 Complete BUSCOs (C)
353 Complete and single-copy BUSCOs (S)
0 Complete and duplicated BUSCOs (D)
7 Fragmented BUSCOs (F)
4 Missing BUSCOs (M)
364 Total BUSCO groups searched

wAlce
C:97.3%[S:61.0%,D:36.3%],F:2.2%,M:0.5%,n:364
354 Complete BUSCOs (C)
222 Complete and single-copy BUSCOs (S)
132 Complete and duplicated BUSCOs (D)
8 Fragmented BUSCOs (F)
2 Missing BUSCOs (M)
364 Total BUSCO groups searched

**Proteobacteria database**

wMeur1
C:68.5%[S:68.5%,D:0.0%],F:2.3%,M:29.2%,n:219
150 Complete BUSCOs (C)
150 Complete and single-copy BUSCOs (S)
0 Complete and duplicated BUSCOs (D)
5 Fragmented BUSCOs (F)
64 Missing BUSCOs (M)
219 Total BUSCO groups searched

wMmer
C:73.5%[S:73.5%,D:0.0%],F:2.3%,M:24.2%,n:219
161 Complete BUSCOs (C)
161 Complete and single-copy BUSCOs (S)
0 Complete and duplicated BUSCOs (D)
5 Fragmented BUSCOs (F)
53 Missing BUSCOs (M)
219 Total BUSCO groups searched

wMeur2
C:83.1%[S:83.1%,D:0.0%],F:1.8%,M:15.1%,n:219
182 Complete BUSCOs (C)
182 Complete and single-copy BUSCOs (S)
0 Complete and duplicated BUSCOs (D)
4 Fragmented BUSCOs (F)
33 Missing BUSCOs (M)
219 Total BUSCO groups searched

wPaur
C:82.2%[S:81.7%,D:0.5%],F:2.7%,M:15.1%,n:219
180 Complete BUSCOs (C)
179 Complete and single-copy BUSCOs (S)
1 Complete and duplicated BUSCOs (D)
6 Fragmented BUSCOs (F)
33 Missing BUSCOs (M)
219 Total BUSCO groups searched

wAlce
C:79.9%[S:51.1%,D:28.8%],F:4.6%,M:15.5%,n:219
175 Complete BUSCOs (C)
112 Complete and single-copy BUSCOs (S)
63 Complete and duplicated BUSCOs (D)
10 Fragmented BUSCOs (F)
34 Missing BUSCOs (M)
219 Total BUSCO groups searched
