## Supplementary table 3 for "Extremely reduced supergroup F *Wolbachia*: transition to obligate insect symbionts"

**Supplementary table 3**: Accessions numbers of the sequences used in phylogenetic and comparative analyses. Blue background = new strains from chewing lice. Green background = taxa included into the phylogenetic analyse (2 gene = fbpA_coxA matrix, multigene = multigene matrix). SG = supergroup. * = assembly done in this study based on the SRA data. MLST = genes retrieved from pubMLST database (see methods).

| | | | | | | Accession number | | | | |
|---|---|---|---|---|---|---|---|---|---|---|
| | | | | | | NCBI | | | MLST | |
| 2 gene | multi gene | Host species | Host taxon | Abbrev. | SG | SRA*/genome assembly | fbpA | coxA | fbpA | coxA |
| | | *Alcedoecus* sp. | louse | *w*Alce | F | SRR5308110* | | | | |
| | | *Menacanthus eurysternus* | louse | *w*Meur1 | F | BioProject: PRJNA768995 | | | | |
| | | *Menacanthus eurysternus* | louse | *w*Meur2 | F | BioProject: PRJNA768995 | | | | |
| | | *Meromenopon meropis* | louse | *w*Mmer | F | SRR8334265* | | | | |
| | | *Penenirmus auritus* | louse | *w*Paur | F | SRR5308137* | | | | |
| | | *Acisoma panorpoides panorpoides* | dragonfly | | F | | KC915265 | KC915236 | | |
| | | *Amblyomma americanum* | tick | | F | | HM061163 | HM061159 | | |
| | | *Anax guttatus* | dragonfly | | F | | KC915285 | KC915256 | | |
| | | *Apilitermes longiceps* | termites | | F | | EF417907 | EF417914 | | |
| | | *Apoica* sp. | wasp | | F | | EU126428 | EU126210 | | |
| | | *Armadillidium vulgare* | carpenter bug | wVulC | B | ALWU00000000 | | | | |
| | | *Atemnus politus* | pseudoscorpion | wAPolK3 | S | JAAXCS000000000 | | | | |
| | | *Atemnus politus* | pseudoscorpion | wAPolK5 | S | WQMQ00000000 | | | | |
| | | *Brachythemis contaminata* | dragonfly | | F | | KC915280 | KC915250 | | |
| | | *Brugia malayi* | filarial nematode | wBm | D | NC_006833 | | | | |
| | | *Carposina sasakii* | moth | wCauA | A | NZ_CP041215 | | | | |
| | | *Cavitermes tuberosus* | termites | | F | | MF953231 | MF953228 | | |
| | | *Chorthippus parallelus* | grasshopper | | F | | JN698879 | JN698878 | | |
| | | *Chrysocoris stolli* | jewel bugs | | F | | | | 410 | 226 |
| | | *Cimex hemipterus* | bed bug | wChem | T | NZ_CP061738 | | | | |
| | | *Cimex lectularius* | bed bug | wCle | F | NZ_AP013028 | | | | |
| | | *Cimex lectularius* | bed bug | wCle | F | | DQ842349 | DQ842275 | | |
| | | *Cruorifilaria tuberocauda* | filarial nematode | wCtub | J | CP046579 | | | | |
| | | *Ctenocephalides felis* | flea | wCfeJ | ? | NZ_CP051157 | | | | |
| | | *Ctenocephalides felis* | flea | wCfeT | ? | NZ_CP051156 | | | | |
| | | *Culex quinquefasciatus* | mosquito | wPip | B | NC_010981 | | | | |
| | | *Dipetalonema caudispina* | filarial nematode | wDcau | J | CP046580 | | | | |
| | | *Diplacodes trivialis* | dragonfly | | F | | KC915276 | KC915246 | | |
| | | *Drosophila melanogaster* | fly | wMel | A | NC_002978 | | | | |
| | | *Drosophila simulans* | fly | wRi | A | NC_012416 | | | | |
| | | *Drosophila simulans* | fly | wNo | B | NC_021084 | | | | |
| | | *Epophthalmia vittata* | dragonfly | | F | | KC915284 | KC915255 | | |
| | | *Folsomia candida* | springtail | wFol | E | NZ_CP015510 | | | | |
| | | *Icosta* sp. | fly | | F | | MF461509 | MF461499 | | |
| | | *Ictinogomphus rapax* | dragonfly | | F | | KC915282 | KC915252 | | |
| | | *Laodelphax striatellus* | planthopper | wStri | B | MUIX00000000 | | | | |
| | | *Madathamugadia hiepei* | filarial nematode | wMhi | F | NZ_WQMP00000000 | | | | |
| | | *Madathamugadia hiepei* | filarial nematode | wMhi | F | | JQ888342 | JQ888307 | | |
| | | *Mansonella (Cutifilaria) perforata* | filarial nematode | | F | | KU255337 | KU255278 | | |
| | | *Melophagus ovinus* | fly | wMelo | F | CACREU02 | | | | |
| | | *Mengenilla moldrzyki* | Strepsiptera | wMen | F | SRX095325* | | | | |
| | | *Nasutitermes ephratae* | termites | | F | | KX036780 | KX024837 | | |
| | | *Nasutitermes itapocuensis* | termites | | F | | KX036782 | KX024839 | | |
| | | *Nasutitermes jaraguae* | termites | | F | | KX036784 | KX024841 | | |
| | | *Neotroponiscus carolii* | Terrestrial isopods | | F | | KX036778 | KX024835 | | |
| | | *Neotroponiscus littoralis* | Terrestrial isopods | | F | | KX036776 | KX024833 | | |
| | | *Nesobasis recava* | damselflies | | F | | MH291053 | MH290892 | | |
| | | *Nesobasis telegastrum* | damselflies | | F | | MH291048 | MH290890 | | |
| | | *Neurothemis tullia* | dragonfly | | F | | KC915283 | KC915254 | | |
| | | *Nilaparvata lugens* | planthopper | wLug | B | MUIY01000000 | | | | |
| | | *Nomada flava* | bee | wNfla | A | NZ_LYUW00000000 | | | | |
| | | *Nycteribia kolenatii* | fly | | F | | MF461503 | MF461494 | | |
| | | *Ocymyrmex picardi* | ant | | F | | EU127822 | EU127606 | | |
| | | *Odontotermes horni* | termites | | F | | GQ422843 | GQ422835 | | |
| | | *Odontotermes* sp. | termites | | F | | GQ422844 | GQ422836 | | |
| | | *Onchocerca ochengi* | filarial nematode | wOo | C | NC_018267 | | | | |
| | | *Onchocerca volvulus* | filarial nematode | wOv | C | NZ_HG810405 | | | | |
| | | *Opistophthalmus capensis* | scorpions | | F | | | | 31 | 30 |
| | | *Opistophthalmus granifrons* | scorpions | | F | | | | 33 | 31 |
| | | *Opistophthalmus litoralis* | scorpions | | F | | | | 57 | 56 |
| | | *Orthetrum glaucum* | dragonfly | | F | | KC915286 | KC915257 | | |
| | | *Osmia caerulescens* | bee | wOc | F | SRR1221705* | KP265901 | | | |
| | | *Paratrechina longicornis* | ant | | F | | | | 226 | 147 |
| | | *Pratylenchus penetrans* | lesion nematodes | wPpe | L | MJMG01000000 | | | | |
| | | *Procornitermes araujoi* | termites | | F | | KX036785 | KX024842 | | |
| | | *Procornitermes lespesii* | termites | | F | | KX036786 | KX024843 | | |
| | | *Pseudolynchia* sp. | fly | | F | | MF461506 | MF461497 | | |
| | | *Supella longipalpa* | cockroach | | F | | | | 414 | 147 |
| | | *Teratodes monticolis* | grasshopper | | F | | | | 125 | 147 |
| | | *Trichogramma pretiosum* | wasp | wTpre | B | NZ_CM003641 | | | | |
| | | *Trithemis pallidinervis* | dragonfly | | F | | KC915267 | KC915237 | | |
| | | *Wuchereria bancrofti* | filarial nematode | wWb | D | NZ_NJBR00000000 | | | | |
| | | *Zelotes* sp. | spider | | F | | | | 475 | 306 |
