## Supplementary table 4 for "Extremely reduced supergroup F *Wolbachia*: transition to obligate insect symbionts"

**Supplementary table 4:** Functional annotations of genes unique for a single genome. Highlighted by blue = new chewing lice strains. The three pantothenate related genes printed in bold blue.

| *w*Meur1 |
|---|
| **3-methyl-2-oxobutanoate_hydroxymethyltransferase__EC_2.1.2.11** |
| **Ketopantoate_reductase_PanG__EC_1.1.1.169** |
| **Pantoate--beta-alanine_ligase__EC_6.3.2.1** |
| 34 hypothetical proteins |
| ***w*Mmer** |
| Uroporphyrinogen_III_decarboxylase__EC_4.1.1.37 |
| 82 hypothetical proteins |
| ***w*Meur2** |
| Proton/glutamate_symport_protein_@_Sodium/glutamate_symport_protein |
| 14 hypothetical proteins |
| ***w*Paur** |
| GTP-binding_protein_Era |
| LSU_ribosomal_protein_L5p__L11e |
| 30 hypothetical proteins |
| ***w*Alce** |
| Zinc_ABC_transporter__periplasmic-binding_protein_ZnuA |
| zinc_protease |
| Arginyl-tRNA_synthetase__EC_6.1.1.19 |
| Malate_dehydrogenase__EC_1.1.1.37 |
| Phospholipase/carboxylesterase_family_protein__EC_3.1.-.- |
| tRNA_nucleotidyltransferase__EC_2.7.7.21___EC_2.7.7.25 |
| Phosphoribosylformylglycinamidine_cyclo-ligase__EC_6.3.3.1 |
| 35 hypothetical proteins |
| ***w*Cle** |
| Hydroxyethylthiazole_kinase__EC_2.7.1.50 |
| Hydroxymethylpyrimidine_phosphate_kinase_ThiD__EC_2.7.4.7 |
| 34 hypothetical protein |
| ***w*Melo** |
| acetyltransferase__GNAT_family |
| 3-oxoacyl-[acyl-carrier_protein]_reductase__EC_1.1.1.100 |
| ATP_synthase_delta_chain__EC_3.6.3.14 |
| Thioredoxin_reductase__EC_1.8.1.9 |
| 2_3_4_5-tetrahydropyridine-2_6-dicarboxylate_N-succinyltransferase__EC_2.3.1.117 |
| Phosphate_transport_system_permease_protein_PstC__TC_3.A.1.7.1 |
| Ferric_iron_ABC_transporter__iron-binding_protein |
| wMelo_-_Lysyl-tRNA_synthetase__class_I___EC_6.1.1.6 |
| D-alanine--D-alanine_ligase__EC_6.3.2.4 |
| DNA_polymerase_III_delta_prime_subunit__EC_2.7.7.7 |
| Uroporphyrinogen-III_synthase__EC_4.2.1.75 |
| rRNA_small_subunit_methyltransferase_H |
| 41 hypothetical proteins |
| ***w*Mhi** |
| Xaa-Pro_aminopeptidase__EC_3.4.11.9 |
| 34 hypothetical proteins |
| ***w*Oc** |
| Glycerol-3-phosphate_dehydrogenase_[NAD_P_+]__EC_1.1.1.94 |
| Cytochrome_bd2__subunit_I |
| putative_Cytochrome_bd2__subunit_II |
| Osmotically_inducible_protein_Y_precursor |
| 38 hypothetical proteins |
