## Supplementary table 5 for "Extremely reduced supergroup F *Wolbachia*: transition to obligate insect symbionts"

**Supplementary table 5:** Overview of metabolic capacities - B vitamines and amino acids

| | | | *w*Meur1 | *w*Mmer | *w*Meur2 | *w*Paur | *w*Alce | *w*Cle | *w*Mhi | *w*Melo | *w*Oc | *w*CfeT | *s*Col |
|---|---|---|---|---|---|---|---|---|---|---|---|---|---|
| B vitamins | Thiamine | iscS | | | | | | | | | | | |
| | | adk | | | | | | | | | | | |
| | | tenA | | | | | | | | | | | |
| | Riboflavin | ribA | | | | | | | | | | | |
| | | ribD | | | | | | | | | | | |
| | | ribB | | | | | | | | | | | |
| | | ribH | | | | | | | | | | | |
| | | ribF | | | | | | | | | | | |
| | | ribE | | | | | | | | | | | |
| | Pyridoxine | pdxJ | | | | | | | | | | | |
| | | pdxH | | | | | | | | | | | |
| | Folate | folA/DHFR | | | | | | | | | | | |
| | | folB | | | | | | | | | | | |
| | | folKP | | | | | | | | | | | |
| | | folC | | | | | | | | | | | |
| | Biotin | bioA | | | | | | | | | | | |
| | | bioB | | | | | | | | | | | |
| | | bioC | | | | | | | | | | | |
| | | bioD | | | | | | | | | | | |
| | | bioF | | | | | | | | | | | |
| | | birA | | | | | | | | | | | |
| | Pantothenate | panB | | | | | | | | | | | |
| | | panG | | | | | | | | | | | |
| | | panC | | | | | | | | | | | |
| | | panD | | | | | | | | | | | |
| amino acids | Asparagine, Aspartate | ansA | | | | | | | | | | | |
| | Arginine | argA | | | | | | | | | | | |
| | | argB | | | | | | | | | | | |
| | | argC | | | | | | | | | | | |
| | | argD | | | | | | | | | | | |
| | | argG | | | | | | | | | | | |
| | | argF (argI, OTC) | | | | | | | | | | | |
| | | argJ | | | | | | | | | | | |
| | | argH | | | | | | | | | | | |
| | Phenylalanine | pheA | | | | | | | | | | | |
| | | aspC | | | | | | | | | | | |
| | Tryptophan | trpD | | | | | | | | | | | |
| | | trpG | | | | | | | | | | | |
| | | trpC | | | | | | | | | | | |
| | | trpA | | | | | | | | | | | |
| | | trpF | | | | | | | | | | | |
| | | trpB | | | | | | | | | | | |
| | Sulphate, cysteine | cysN | | | | | | | | | | | |
| | | cysD | | | | | | | | | | | |
| | | cysC | | | | | | | | | | | |
| | | cysH | | | | | | | | | | | |
| | | cysI | | | | | | | | | | | |
| | | cysJ | | | | | | | | | | | |
| | | cysQ | | | | | | | | | | | |
| | | cysE | | | | | | | | | | | |
| | | cysK | | | | | | | | | | | |
| | Methionine | metA | | | | | | | | | | | |
| | | metB | | | | | | | | | | | |
| | | metC | | | | | | | | | | | |
| | | metE | | | | | | | | | | | |
| | Lysine | asd | | | | | | | | | | | |
| | | dapA | | | | | | | | | | | |
| | | dapB | | | | | | | | | | | |
| | | dapD | | | | | | | | | | | |
| | | dapE | | | | | | | | | | | |
| | | dapF | | | | | | | | | | | |
| | | lysA | | | | | | | | | | | |
| | | thrA | | | | | | | | | | | |
| | Threonine | thrB | | | | | | | | | | | |
| | | thrC | | | | | | | | | | | |
| | Leucine | leuA | | | | | | | | | | | |
| | | leuC | | | | | | | | | | | |
| | | leuD | | | | | | | | | | | |
| | | leuB | | | | | | | | | | | |
| | Glycine | glyA | | | | | | | | | | | |
| | Histidine | hisA | | | | | | | | | | | |
| | | hisB | | | | | | | | | | | |
| | | hisC | | | | | | | | | | | |
| | | hisD | | | | | | | | | | | |
| | | hisF | | | | | | | | | | | |
| | | hisG | | | | | | | | | | | |
| | | hisH | | | | | | | | | | | |
| | | hisI | | | | | | | | | | | |
| | Serine | serC | | | | | | | | | | | |

| present | absent |
|---|---|

**Supplementary table 5:** Overview of metabolic capacities - secretion systems and ABC transporters

| | | *w*Meur1 | *w*Mmer | *w*Meur2 | *w*Paur | *w*Alce | *w*Cle | *w*Mhi | *w*Melo | *w*Oc | *w*CfeT | *s*Col |
|---|---|---|---|---|---|---|---|---|---|---|---|---|
| **secretion system** | **Type I - tolC** | present | present | present | present | present | present | present | present | present | present | absent |
| | **Type II - gspD** | absent | absent | absent | absent | absent | absent | absent | absent | absent | absent | absent |
| | **Sec-SRP - secA** | present | present | absent | present | present | present | present | present | present | present | present |
| | **Sec-SRP - secB** | present | present | present | present | present | present | present | present | present | present | absent |
| | **Sec-SRP - secD** | present | present | present | present | present | present | present | present | present | present | present |
| | **Sec-SRP - secF** | present | present | present | present | present | present | present | present | present | present | present |
| | **Sec-SRP - secG** | present | present | present | present | present | present | present | present | present | present | present |
| | **Sec-SRP - secY** | present | present | present | present | present | present | present | present | present | present | present |
| | **Sec-SRP - YajC** | present | present | present | present | present | present | present | present | present | present | present |
| | **Sec-SRP - YidC** | present | present | present | present | present | present | present | present | present | present | present |
| | **Sec-SRP - ftsY** | present | present | present | present | present | present | present | present | present | present | absent |
| | **Sec-SRP - ffh** | absent | absent | absent | absent | absent | absent | absent | absent | absent | absent | absent |
| | **secE** | present | present | present | present | present | present | present | present | present | absent | present |
| | **Twin arginine targeting - TatA** | present | present | present | present | present | present | present | present | present | absent | absent |
| | **Twin arginine targeting - TatC** | present | present | present | present | present | present | present | present | present | present | absent |
| | **Type IV - virB10** | present | present | present | present | present | present | present | present | present | present | absent |
| | **virB3** | present | present | present | present | present | present | present | present | present | absent | absent |
| | **virB4** | present | present | present | present | present | present | present | present | present | present | absent |
| | **virB9** | present | present | present | present | present | present | present | present | present | present | absent |
| | **virB6** | present | present | present | present | present | present | present | present | present | present | absent |
| | **virB8** | present | present | present | present | present | present | present | present | present | present | absent |
| | **virB11** | present | present | present | present | present | present | present | present | present | present | absent |
| | **virD4** | present | present | present | present | present | present | present | present | present | present | absent |
| **ABC transporters** | **CcmA** | present | present | present | present | present | present | present | present | present | present | absent |
| | **CcmB** | present | present | present | present | present | present | present | present | present | present | absent |
| | **CcmC** | present | present | present | present | present | present | present | present | present | present | absent |
| | **haem exporter - PstA** | present | present | present | absent | present | present | present | present | present | present | absent |
| | **haem exporter - PstB** | present | present | present | present | present | present | present | present | present | present | absent |
| | **haem exporter - PstC** | present | present | present | present | present | present | present | present | present | present | absent |
| | **haem exporter - PstS** | present | present | present | present | present | present | present | present | present | present | absent |
| | **phosphate trasport system - lolC_E** | present | present | present | present | present | present | present | present | present | present | present |
| | **phosphate trasport system - lolD** | present | present | present | present | present | present | present | present | present | present | present |
| | **lioprotein releasing systme - ZnuA** | present | present | present | present | present | present | present | present | present | present | absent |
| | **lioprotein releasing systme - ZnuB** | present | present | present | present | present | present | present | present | present | present | absent |
| | **lioprotein releasing systme - ZnuC** | present | present | present | present | present | present | present | present | present | present | absent |
| | **zinc transport system - BioY** | absent | absent | present | present | present | present | present | present | present | absent | absent |
| | **biotin trasport system - AfuA** | absent | absent | present | present | present | present | present | present | present | present | absent |
| | **biotin trasport system - AfuB** | absent | absent | present | present | present | present | present | present | present | present | absent |
| | **biotin trasport system - AfuC** | absent | absent | present | present | present | present | present | present | present | present | absent |
| | **phospholipid trasport system - MlaC** | absent | present | present | present | present | present | present | present | present | present | present |
| | **phospholipid trasport system - MlaD** | absent | absent | present | present | present | present | present | present | present | absent | present |
| | **phospholipid trasport system - MlaE** | absent | absent | present | present | present | present | present | present | present | present | present |
| | **phospholipid trasport system - MlaF** | absent | absent | present | present | present | present | present | present | present | present | present |

| | |
|---|---|
| **present** | **absent** |

**Supplementary table 5:** Overview of metabolic capacities - cellular processes

| | |
|---|---|
| present | absent |

| Category | Gene | *w*Meur1 | *w*Mmer | *w*Meur2 | *w*Paur | *w*Alce | *w*Cle | *w*Mhi | *w*Melo | *w*Oc | *w*CfeT | *s*Col |
|---|---|---|---|---|---|---|---|---|---|---|---|---|
| Fatty acids | fabK | present | present | present | present | present | present | present | present | present | present | present |
| | fabF | present | present | present | present | present | present | present | present | present | present | absent |
| | fabG | present | present | present | present | present | present | present | present | present | present | present |
| | fabZ | present | present | present | present | present | present | present | present | present | present | present |
| | fabI | present | present | present | present | present | present | present | present | present | present | present |
| | fabD | present | present | present | present | present | present | present | present | present | present | present |
| | fabH | present | present | present | present | present | present | present | present | present | present | present |
| Purines | prsA | absent | absent | absent | absent | absent | absent | absent | absent | absent | absent | absent |
| | purB | present | present | present | present | present | present | present | present | present | present | present |
| | purC | present | present | present | present | present | present | present | present | present | present | absent |
| | purD | present | present | present | present | present | present | present | present | present | present | absent |
| | purF | present | present | present | present | present | present | present | present | present | present | absent |
| | purK | present | present | present | present | present | present | absent | present | present | present | absent |
| | purN | present | present | present | present | present | present | present | present | present | present | absent |
| | purQ | present | present | present | present | present | present | present | present | present | present | absent |
| | purSL | present | present | present | present | present | present | present | present | present | present | absent |
| | purM | present | present | present | present | present | present | present | present | present | present | absent |
| | purE | present | present | present | present | present | present | present | present | present | present | absent |
| | purA | present | present | present | present | present | present | present | present | present | present | present |
| | purH | present | present | present | present | present | present | present | present | present | present | absent |
| | gmk | absent | absent | absent | absent | absent | absent | absent | absent | absent | absent | absent |
| | dgt | absent | present | present | present | present | present | present | present | present | present | absent |
| | surE | present | present | present | present | present | present | present | present | present | present | absent |
| | nrdA | absent | absent | absent | absent | absent | absent | absent | absent | absent | absent | absent |
| | nrdB | absent | absent | absent | absent | absent | absent | absent | absent | absent | absent | absent |
| | guaA | present | present | present | present | present | present | present | present | present | present | present |
| | guaB | absent | absent | absent | absent | absent | absent | absent | absent | absent | absent | absent |
| Pyrimidines | carB | present | present | present | present | present | present | present | present | present | present | absent |
| | pyrB | present | present | present | present | present | present | present | present | present | present | absent |
| | pyrC | absent | absent | absent | absent | absent | absent | absent | absent | absent | absent | absent |
| | pyrD | absent | absent | absent | absent | absent | absent | absent | absent | absent | absent | absent |
| | pyrE | present | present | present | present | present | present | present | present | present | present | absent |
| | pyrF | present | present | present | present | present | present | absent | absent | present | present | absent |
| | pyrH | present | present | present | present | present | present | present | present | present | present | present |
| | pyrG | present | present | present | present | present | present | present | present | present | present | present |
| | surE | present | present | present | present | present | present | present | present | present | present | absent |
| | ndk | present | present | present | present | present | present | present | present | present | present | absent |
| | dut | present | present | present | present | present | present | present | present | present | present | present |
| | tmk | present | present | present | present | present | present | present | present | absent | present | present |
| | thyX | present | present | present | present | present | present | present | present | present | present | absent |
| | carA | present | present | present | present | present | present | present | present | present | present | absent |
| Porphyrin | gltX | absent | absent | absent | absent | absent | absent | absent | absent | absent | absent | absent |
| | hemB | absent | present | present | present | present | present | present | present | present | present | absent |
| | hemC | absent | present | present | present | present | present | present | present | present | present | absent |
| | hemD | absent | present | present | present | present | present | present | present | present | present | absent |
| | hemE | absent | present | present | present | present | present | present | present | present | present | absent |
| | hemF | absent | absent | absent | absent | absent | absent | absent | absent | absent | absent | absent |
| | hemJ | absent | present | present | present | present | present | present | present | present | present | absent |
| | ctaB | absent | absent | absent | absent | absent | absent | absent | absent | absent | absent | absent |
| | ctaA | absent | absent | absent | absent | absent | absent | absent | absent | absent | absent | absent |
| | bfr | present | present | present | present | present | present | present | present | present | present | absent |
| Oxidative phosphorylation | RIP1 | absent | absent | absent | absent | absent | absent | absent | absent | absent | absent | absent |
| | CYTB | present | present | present | present | present | present | present | present | present | present | absent |
| | CYC1 | present | present | present | present | present | present | present | present | present | present | absent |
| | COX10 | present | present | present | present | present | present | present | present | present | present | present |
| | COX11 | present | present | present | present | present | present | present | present | present | present | absent |
| | COX15 | present | present | present | present | present | present | present | present | present | present | absent |
| | CoxC | present | present | present | present | present | present | present | present | present | present | absent |
| | CoxA | present | present | present | present | present | present | present | present | present | present | absent |
| | CoxB | present | present | present | present | present | present | present | present | present | present | absent |
| | Cyt. bd ubiquinol oxidase - cydA | absent | absent | present | present | absent | absent | absent | present | present | absent | absent |
| | Cyt. bd ubiquinol oxidase - cydB | absent | absent | present | present | absent | absent | absent | present | present | absent | absent |
| Cell cycle | murG | present | present | present | present | present | present | present | present | present | present | present |
| | rseP | present | present | present | present | present | present | present | present | present | present | absent |
| | lon | present | present | present | present | present | present | present | present | present | present | present |
| | dnaB | present | present | present | present | present | present | present | present | present | present | present |
| | dnaA | present | present | present | present | present | present | present | present | present | present | absent |
| | PerP | absent | absent | absent | absent | absent | absent | absent | absent | absent | absent | present |
| | cell division - ftsW | absent | present | present | present | present | present | present | present | present | present | present |
| | cell division - ftsZ | present | present | present | present | present | present | absent | present | present | present | present |
| | cell division - ftsQ | present | present | present | present | present | present | present | present | present | present | absent |
| | cell division - ftsA | present | present | present | present | present | present | present | present | present | present | present |
| | two-component - pleC | absent | present | present | present | present | present | present | present | present | present | absent |
| | two-component - pleD | absent | present | present | present | present | present | present | present | present | present | absent |
| | ATP-dependent Clp - clpX | present | present | present | present | present | present | present | present | present | present | present |
| | ATP-dependent Clp - clpP | present | present | present | present | present | present | present | present | present | present | present |
| Recombination | ssb | present | present | present | present | present | present | present | present | present | present | present |
| | polA | present | present | present | present | present | present | present | present | present | present | present |
| | dnaE | present | present | present | present | present | present | present | present | present | present | present |
| | recJ | present | present | present | present | present | present | present | present | present | present | absent |
| | recA | absent | absent | present | present | present | present | present | present | present | present | present |
| | recF | absent | absent | present | present | absent | present | present | present | present | present | absent |
| | recO | absent | absent | present | present | absent | present | present | absent | present | present | present |
| | recR | absent | absent | present | present | present | present | present | present | present | present | present |
| | recG | absent | absent | present | present | present | present | present | present | present | present | present |
| | ruvA | absent | absent | present | present | present | present | present | present | present | present | present |
| | ruvB | absent | absent | present | present | present | present | present | present | present | present | present |
| | ruvC | absent | absent | present | present | present | present | present | present | present | present | present |
| Glycerophospholipids | gpsA | present | present | present | present | present | present | present | present | present | present | present |
| | plsY | present | present | present | present | present | present | present | present | present | present | present |
| | plsC | present | present | present | present | present | present | present | present | present | present | present |
| | pssA | present | present | present | present | present | present | present | present | present | present | present |
| | psd | present | present | present | present | present | present | present | present | present | present | present |
| | cdsA | absent | absent | absent | absent | absent | absent | absent | absent | absent | absent | absent |
| | pgpA | present | present | present | present | present | present | present | present | absent | absent | present |
| | pgsA | absent | absent | absent | absent | absent | absent | absent | absent | absent | absent | present |
| | araM | absent | present | present | present | present | present | present | present | present | absent | absent |
| | dgkA | absent | absent | absent | absent | absent | absent | absent | absent | absent | present | absent |
| | pld | absent | absent | absent | absent | absent | absent | absent | absent | absent | absent | absent |
